# Exosome secretion is required for sonic hedgehog dispersal and signal gradient formation in the embryonic limb mesenchyme

**DOI:** 10.64898/2025.12.05.692671

**Authors:** Sean Corcoran, Joshua Fisher, Timothy A. Sanders, Edwin Munro

**Author notes:** **Contact info:** Edwin Munro Molecular Genetics and Cell Biology University of Chicago 920 E 58th St CLSC 901.

## Abstract

Carrier-assisted diffusion and cytoneme transport have been postulated to disperse Hedgehog across diverse embryonic tissues, yet their relative contributions to patterning mesenchymal tissues remains poorly understood. Here, we combine novel signaling assays with quantitative microscopy to establish exosome secretion as a predominant and adaptable mechanism for Sonic Hedgehog (Shh) dispersal. Introducing a novel synchronous release system to visualize Shh trafficking in embryonic tissues, we demonstrate that Shh traffics through the exosome biogenesis pathway in the limb bud mesenchyme. Shh-bound exosomes diffuse through extracellular space, and can also bind and travel along cytonemes, providing a potential mechanism for directed and/or long-range transport. Using a synthetic patterning assay, we show that exosome secretion is essential to establish short-range Shh gradients *in vitro*. We propose that exosome-based Shh secretion, combined with different modes of extracellular transport, provides a tunable mechanism to sculpt Shh gradients on different length and time scales, across different embryonic tissues.

**HIGHLIGHTS:**

- A novel synchronous release system reveals trafficking dynamics in embryonic cells
- Sonic hedgehog is packaged and secreted on exosomes for extracellular dispersal
- Exosome secretion is required to establish short-range Hedgehog gradients
- Diffusion and cytoneme transport provide tunable exosome dispersal strategies

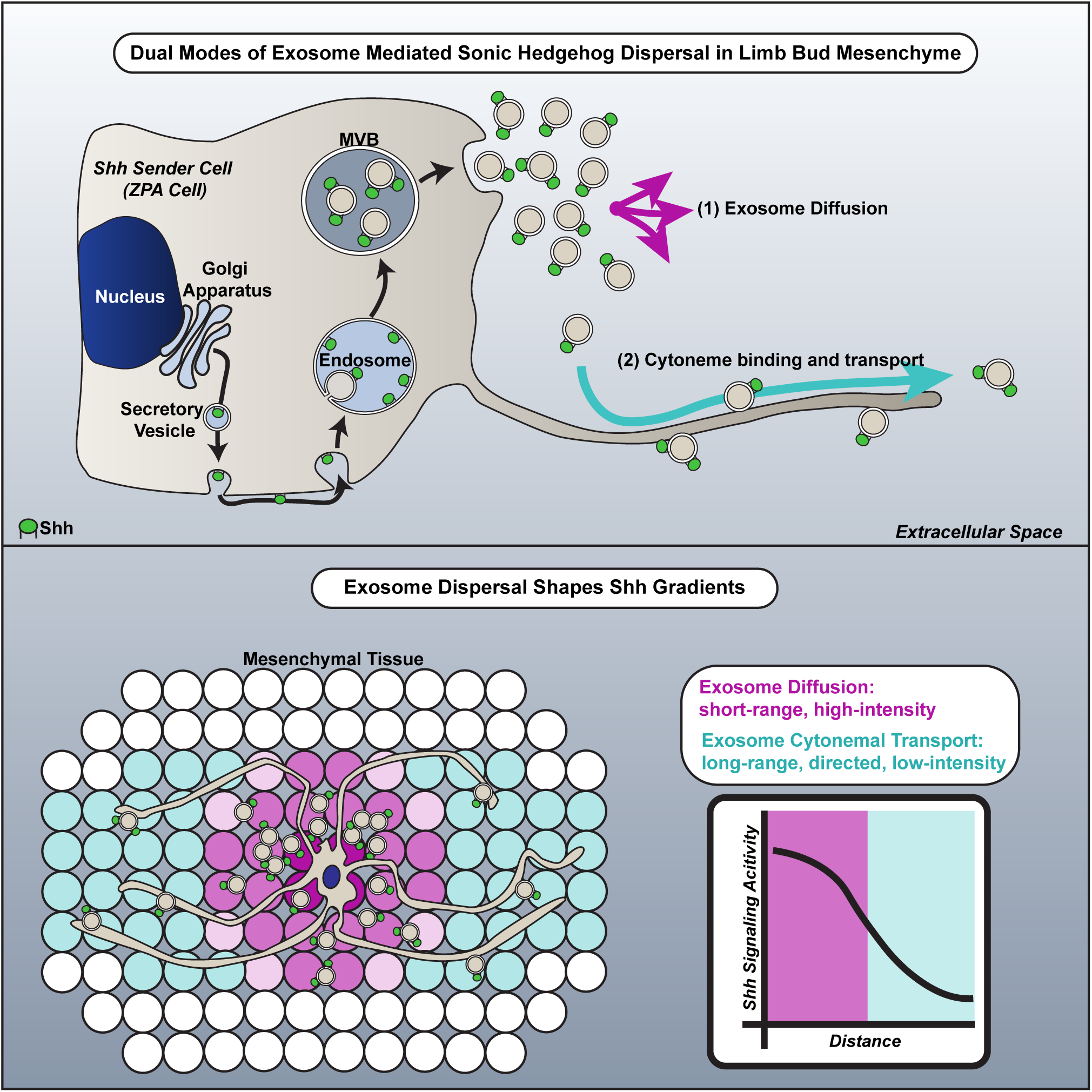

## INTRODUCTION

Embryos use a small number of highly conserved developmental signaling molecules, known as morphogens, to pattern cell fates across diverse tissues. Morphogens are secreted by source cells and dispersed across target tissues to shape gradients of signaling activities that are interpreted by target cells. How the same morphogens are dispersed in different ways to shape gradients across tissues with very different architectures (e.g. epithelial vs mesenchymal) and sizes remains poorly understood.

Hedgehog (Hh) signaling is a broadly conserved pathway which controls many different cellular behaviors during development and homeostasis, including cell fate decisions, proliferation, survival, migration, and stem cell maintenance.^1^ Hh morphogens are secreted by source cells and dispersed across tissues where they bind to the Patched (PTCH) receptor on target cells to relieve inhibition of the receptor-like signal transducer Smoothened (SMO). Active SMO, in turn, promotes the stabilization and nuclear translocation of active Glioma Associated Oncogene (GLI) transcription factors to control transcription of Hh target genes. Among the three vertebrate Hh paralogs, Sonic Hedgehog (Shh) is the most broadly expressed and the best studied, with well-established roles in patterning epithelial and mesenchymal tissues including the neural tube, face, and the developing limb bud.^2–4^ During its biosynthesis, Shh gains two lipid modifications: a palmitate at its amino terminus and cholesterol at its carboxy terminus. Both of these modifications are required for proper spatial distribution and bioactivity of Shh.^5–8^ However, they also confer strong hydrophobicity that tethers Shh to the producing cell’s plasma membrane requiring the transporter-like protein Dispatched (DISP) to mediate release.^9–11^ Once released, these lipid moieties restrict the ability of Shh to diffuse freely through the aqueous extracellular space. Thus, additional mechanisms are required to overcome these restrictions and promote the dispersal of Shh across target tissues. To date, two major classes of mechanisms have been proposed: carrier-assisted diffusion and active transport.

One class of mechanisms involves molecular-scale interactions that facilitate diffusion through extracellular (EC) space. Multimerization of Shh, transfer to high density lipoproteins (HDL), and association with Scube proteins all can promote carrier-assisted diffusion by shielding the hydrophobic moieties of Shh in the EC space.^12–14^ For example, recent studies showed that addition of Scube1 in media greatly increases both the amplitude and the length scale of Shh signaling gradients *in vitro*.^15^

Recent work highlights another important mode of carrier-assisted diffusion involving the packaging of Shh/Hh into small secreted extracellular vesicles. Small extracellular vesicles (sEVs) are secreted by nearly all eukaryotic cells and function to transport lipids, nucleic acids, and proteins from cell to cell.^16^ There are two key classes of sEVs: 1) exosomes and 2) ectosomes. Exosomes are smaller, 30-150 nm in diameter, and their generation is endocytosis dependent. Specifically, they form via endocytosis of material from the plasma membrane (PM), invagination or inward budding of endosomal membrane to form a multivesicular body (MVB) containing intralumenal vesicles, and eventual fusion of the MVB with the PM to release the intralumenal vesicles into the extracellular space as “exosomes”.^17,18^ Ectosomes are larger vesicles, 100-1000 nm in diameter, that are generated by the outward budding of PM and eventual shedding of vesicles to the extracellular space.^19^ Although exosomes and ectosomes contain some of the same molecules, they can be distinguished from one another by their mode of biogenesis. Previous work has shown that Hh is associated with bioactive sEVs and that Shh+ sEVs are bioactive.^20^ Moreover, inhibition of endocytosis can reduce both Hh secretion and the spatial extent of epithelial Hh signaling gradients, suggesting that these sEVs may be exosomes.^21–23^

In addition to carrier-assisted diffusion, a growing body of work suggests that cells harness specialized signaling filopodia known as cytonemes to transport Hh across tissues. Hh ligands and receptor/co-receptors have been shown to travel along these structures in the chick limb bud mesenchyme, Drosophila imaginal disc epithelia, mouse neural tube, and in cell culture models.^24–27^ Inhibiting cytonemes reduced signaling gradients in the Drosophila wing disc epithelium, abdominal epidermis, and in the mouse neural tube epithelium but had no effect on signaling gradients in fibroblast cells cultured *in vitro*.^15,27,28^ Interestingly, several recent papers described the association of Hh positive exovesicles with cytonemes in Drosophila epithelia and in NIH-3T3 fibroblasts, hinting at more complicated relationships between carrier-assisted diffusion and direct transport of Hh proteins.^23,26^

In summary, a variety of different mechanisms have been proposed to shape Hh dispersal *in vivo*. However, the bulk of work thus far has been performed in epithelial tissues which are formed by tightly packed, polarized cells organized into sheets. However, mesenchymal tissues, such as cartilage, bone, and connective tissue, represent a large fraction of the vertebrate body plan and are characterized by loosely packed apolar cells embedded in a complex extracellular matrix. Although Shh signaling is known to mediate the patterning of many mesenchymal tissues, including the developing face and limb,^3,4^ the mechanisms that govern Shh dispersal in these tissues remains less clear. Although immortalized cells *in vitro* can model some aspects Shh dispersal from mesenchyme-like cells and are accessible to biochemistry and high-resolution imaging, they lack much of the endogenous machinery that mediates Shh signaling *in vivo* and thus the developmental relevance of these studies remains unclear. Thus, there continues to be a key gap in understanding how Shh transport works in mesenchymal tissues.

To overcome this gap in understanding, we investigated the mechanisms of Shh dispersal in the chicken embryonic limb bud mesenchyme, a classic model for Shh-mediated tissue patterning. Within the limb bud mesenchyme, a posterior region, known as the zone of polarizing activity (ZPA), is a Shh-expressing signaling center that patterns the anterior-posterior axis of the limb bud mesenchyme and indirectly promotes limb outgrowth via a feedback loop with Fibroblast Growth Factor (FGF) signaling.^4,29,30^ However, the mechanisms that shape gradients of Shh signaling away from the ZPA remain poorly understood, Individual loss of function and knockout mutants of Scube1, Scube2, and Scube3 in mice show normal number and identity of digits in the limb suggesting that Scube proteins may be dispensable or act redundantly for limb patterning *in vivo*.^31–34^ However, roles for other modes of carrier-assisted diffusion have not been explored. Shh has been shown to travel along cytonemes made by limb bud mesenchyme cells *in vivo*, but what regulates cytonemal transport of Shh, and how it contributes to pattering the limb bud, remain unknown.^24^

To address these questions, we introduce cultured primary limb bud mesenchymal cells (PLMCs) as a tractable model system for functional analysis of Shh trafficking and dispersal. We deploy a novel synchronous release system that allows us to visualize Shh trafficking with high spatial and temporal resolution in cultured PLMCs and to confirm our observations *in vivo*. We show that a large fraction of Shh is packaged into exosomes for release into extracellular space, identifying exosome-based secretion as a dominant pathway for Shh dispersal. We find that exosomes disperse both by diffusion and through transport along cytonemes. We show that exosome release, but not cytoneme-based transport, is required to form short-range signaling gradients in a synthetic tissue composed of PLMCs cultured on a monolayer of Shh responder fibroblasts. However, Shh-bound secreted exosomes travel along the longer, dynamic cytonemes observed *in vivo*, pointing to a larger role for cytonemes in promoting longer-range, directed transport during normal limb development. Overall, our study establishes exosome-mediated Shh dispersal as a fundamental, and versatile basis for Shh signaling and mesenchymal tissue patterning.

## RESULTS

### Primary Limb Mesenchymal Cells are a live imaging conducive model system to study Shh dispersal dynamics

Recent studies of chick limb bud mesenchyme cells emphasize potential roles for cytoneme-based transport in Shh dispersal.^24^ However, efforts to assess the relative contributions of these and other modes of transport to Shh dispersal *in vivo* have been limited by the difficulty of achieving high resolution imaging of relevant dynamics. While cultured immortalized cells are better suited for high resolution live imaging, they lack the endogenous dispersal machinery expressed by mesenchymal cells *in vivo.* To bridge this gap, we established primary cultures of Limb Bud Mesenchymal Cells (PLMCs) as a tractable *in vitro* system to study Shh dispersal dynamics with direct relevance to limb bud patterning *in vivo*. We dissected Hamburger and Hamilton (HH) stage 22 embryonic chick limb buds, dissociated them into single cells, electroporated the cells with plasmids of interest, and then plated them at confluency to generate a pseudo 3D (mono to bilayer) culture (Figure 1A, detailed in Methods).

**Figure 1:**
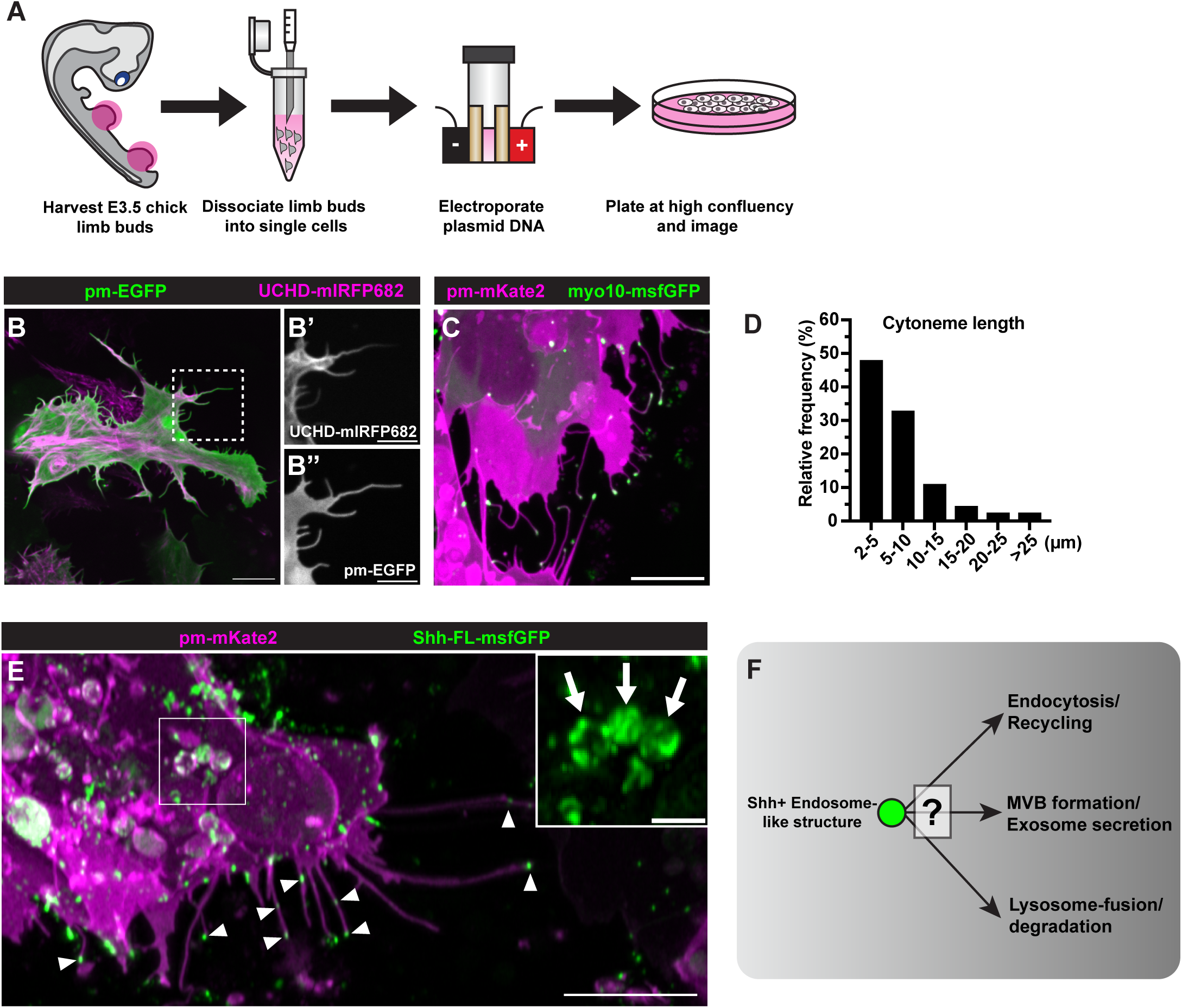
Primary Limb Mesenchymal Cell (PLMC) culture provides a tractable model to study Shh trafficking and dispersal dynamics. A) Schematic of PLMC generation protocol. B) A PLMC electroporated with plasmids encoding palmitoylated (pm)-EGFP (membrane) and mIRFP682 tagged Utrophin-Calponin Homology Domain (F-Actin) - Scale Bar = 10μm. White box marks magnified view displaying protrusions both labeled with F-Actin (B’) and membrane (B’’) - Scale Bar = 5μm. C) A PLMC electroporated with plasmids encoding pm mKate2 (membrane) and msfGFP tagged myosin10 HMM - Scale Bar = 10μm. D) Histogram displaying the relative frequency of the lengths of all cytonemes over 2 microns. E) Micrograph depicting PLMC electroporated with pm-Kate2 to mark cell membrane and Shh-FL-msfGFP. Rolling ball (20 pixel) background subtraction was performed for enhanced visualization of subcellular structures. Shh is decorating cytonemes (arrowheads) - Scale Bar = 10μm. Additionally, the white box and inset depict large, spherical Shh+ endosome-like structures in the cell soma (arrows) - Scale Bar = 2μm. F) Diagram highlighting the possible identities/fates of Shh+ endosome-like structures.

We first sought to verify that PLMCs display core elements of the Shh transport machinery observed in limb bud mesenchyme *in vivo*. Using palmitoylated-EGFP (pm-EGFP) and mIRFP682 tagged Utrophin-Calponin Homology Domain (UCHD-mIRFP682) to label plasma membranes and filamentous actin (F-actin) respectively, we observed that cultured PLMCs project numerous F-actin based filopodia that resemble the cytonemes observed *in vivo* (Figure 1B-B’’), although they are shorter in length (Figure 1D). As observed *in vivo*, these cytonemes undergo dynamic cycles of extension and retraction, and they accumulate the motor domain of the atypical motor Myosin10 (Myo10-HMM-msfGFP) at their tips (Figure 1C, Movie 1). Finally, we observed directed transport of full length Shh tagged with msfGFP (Shh-FL-msfGFP) along the cytonemes of PLMCs, as previously reported for limb bud mesenchyme cells *in vivo* (Figure 1E and Movie 2).

While exogenous expression of Shh can induce cytoneme formation in immortalized cell culture lines^26^, the length and number of cytonemes did not vary with expression levels of exogenous Shh-FL-msfGFP in cultured PLMCs (Supplemental Figure 1). Thus, as *in vivo*, cytoneme production is independent of Shh expression in primary limb bud mesenchyme cells. Altogether, these results show that cultured PLMCs retain many of the known characteristics of limb mesenchymal cells *in vivo*, thus providing an ideal model to study the mechanisms guiding intercellular Shh dispersal.

### Shh traffics to both cytonemes and endosome-like structures rapidly after synthesis

In addition to co-localizing with cytonemes, Shh-FL-msfGFP accumulates within large, spherical endosome-like structures in the cell body (Figure 1E, inset). In principle, these Shh+ endosome-like structures could reflect pools of Shh undergoing recycling, degradation, or packaging into multivesicular bodies (MVBs) for exosome secretion. To distinguish these possibilities, we developed methods to visualize and interrogate secretory trafficking in Limb Bud Cells. One common method, known as Retention Using Selective Hooks (RUSH),^35^ uses high affinity binding of streptavidin binding protein (SBP) to streptavidin to trap secretory cargo in the endoplasmic reticulum (ER). Upon biotin addition, the cargo is released from the ER and its passage through the secretory pathway can be visualized. Unfortunately, our attempts to implement RUSH in limb bud mesenchymal cells were hampered by the high levels of endogenous biotin observed in the chicken yolk/embryos, which prevent the constitutive retention of cargo (Supplemental Figure 2C, 2D bottom panel).^36^ Importantly, many vertebrate embryos, and primary cell lines derived therefrom, express high levels of endogenous biotin, and thus leaky retention of cargo is likely to limit the use of RUSH in these contexts as well.

To overcome this limitation, we designed a functionally analogous trap and release system using Chemically Disruptable Heterodimer (CDH) protein off switches to construct a releasable protein trap. CDH switches consist of a Bcl-xL receptor and a synthetic CDH binder (CB), deemed LD3 in the original publications, that bind one another with high affinity, but dissociate upon addition of a small molecule inhibitor of Bcl-xL (A1155463, hereon A1).^37,38^ To create the ER trap for our CDH synchronous release system, we added the ER localization sequence from the STIM1 gene and the ER retention signal (KDEL) to the Bcl-xL receptor. To make a trappable form of Shh, we inserted the CB moiety upstream of the msfGFP into our full length, msfGFP tagged Shh (Figure 2B). When both constructs are expressed in the same cell, Shh-CB-sfGFP should be trapped in the ER until the addition of A1 (Figure 2A). We systematically evaluated a panel of point-mutated variants of the CB moiety, with different affinities for the Bcl-xL receptor,^38^ and determined that the Shh-CB(D138A) variant is very tightly trapped in the ER in both PLMCs and *in vivo,* yet is rapidly and efficiently released upon exposure to A1 (Supplemental Figure 2A-B, 2D bottom panels). Thus, we used the CB(D138A) moiety (hereon CB) for all subsequent experiments.

**Figure 2:**
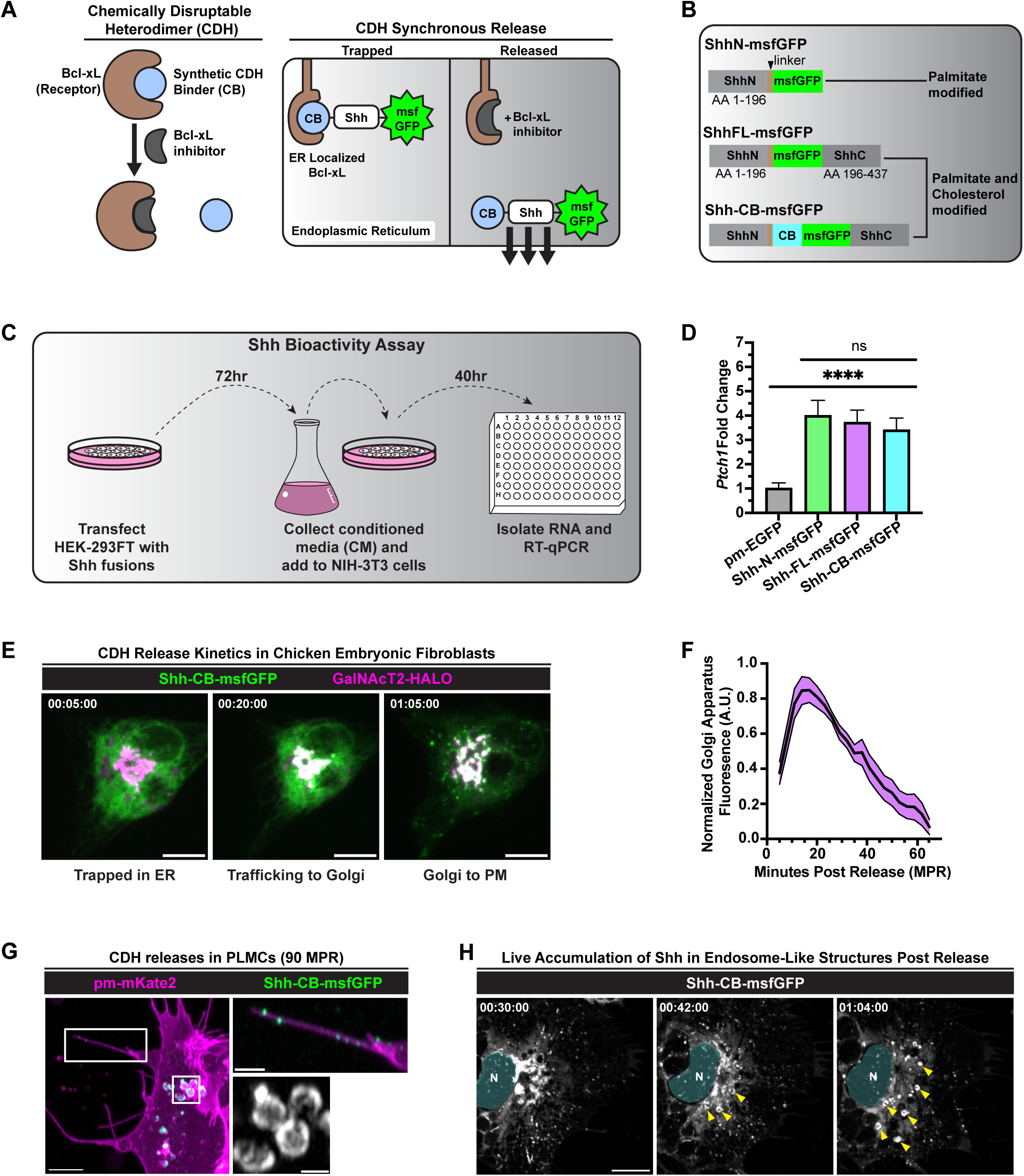
A novel CDH synchronous release system reveals that Shh rapidly accumulates in endosome-like structures after leaving the Golgi Apparatus. A) Schematic of Chemically Disruptable Heterodimer (CDH) protein switch and novel CDH synchronous release system. B) Diagram representing the design of the fusion proteins that allows for normal processing with both palmitate and cholesterol addition. C) Diagram depicting conditioned media (CM) bioactivity assay. D) RT-qPCR shows high levels of Ptch1 upregulation in all fusion protein CM conditions (n = 3 biological replicates, +/- SD, One-way ANOVA with Tukey’s Multiple Comparisons performed). E) Stills from timelapse of CDH release of Shh-CB-msfGFP from the ER in Chicken Embryonic Fibroblasts (CEFs), note the Golgi apparatus marker GalNAcT2-HaloTag (magenta) was co-expressed – Scale Bar = 10 μm. F) Depicts a kinetics graph that measured the normalized Shh-CB-msfGFP fluorescence in the Golgi over the timescale of the CDH release (n = 12 cells/releases) (Mean trace +/- SEM). Shh peaks in the Golgi around 20 MPR and then reduces indicating its trafficking through the rest of secretory pathway. G) Micrograph of Airyscan Super Resolution imaged PLMC cell 90 MPR – Scale Bar = 5 μm. Rolling ball (15 pixel) background subtraction was performed for enhanced visualization of subcellular structures. Top white box shows magnified and brightened view of Shh localized to a cytoneme – Scale Bar = 2 μm. Bottom white box shows magnified view of Shh+ endosome-like structures – Scale Bar = 1 μm. Note inside the lumen of the endosome-like structure exists smaller intralumenal vesicles. H) Timelapse of CDH release of Shh in PLMCs starting 30mpr - Scale Bar = 10μm. Rolling ball (50 pixel) background subtraction was performed for enhanced visualization of subcellular structures. Around 42 MPR, Shh begins accumulating in spherical, endosomal like structures (yellow arrowheads). Nucleus was pseudo colored cyan.

To confirm that the Shh-CB-msfGFP fusion protein can still be secreted and retains normal bioactivity, we collected conditioned media from HEK-293FT cells transfected with Shh-CB-msfGFP, Shh-N-msfGFP (non-cholesterol modified), Shh-FL-msfGFP (cholesterol modified), or pm-EGFP as a control. Then we incubated Shh-responsive NIH-3T3 cells with conditioned media for 40 hours and performed RT-qPCR against the Shh target gene *Ptch1* (Figure 2C). We observed that conditioned media from cells expressing all three Shh variants upregulated *Ptch1* expression compared to the pm-EGFP control, suggesting that Shh-CB-msfGFP can both be properly secreted and remains biologically active (Figure 2D).

To characterize the kinetics of Shh release and post-release trafficking, we expressed the ER-localized trap, Shh-CB-msfGFP, and a Golgi apparatus (GA) marker GalNAcT2-HaloTag (GalNAcT2-HALO) in chicken embryonic fibroblasts. We added A1 and visualized release in live cells via spinning disk microscopy (Figure 2E, Movie 3). After 5 minutes of incubation with A1, Shh-CB-msfGFP remained tightly trapped in the ER, suggesting that the CDH trap does not leak (Figure 2E, left panel). By 20 minutes post release (MPR), Shh-CB-msfGFP began to colocalize in the Golgi apparatus indicated by the overlap with the GalNacT2-Halo signal (Figure 2E, middle panel). By 65 MPR, Shh-CB-msfGFP could be seen as punctate structures moving through the cell body suggesting that Shh can fold properly and move through the secretory system efficiently (Figure 2E, right panel, Movie 3). Quantifying GA-associated Shh-CB-msfGFP signal vs time confirmed that Shh releases with similar kinetics across multiple cells (Figure 2F). These data confirm that the CDH system provides a reproducible and efficient way to follow secretory trafficking of Shh.

Having validated the CDH trap and release system, we turned to analyzing Shh trafficking dynamics in limb bud cells. We expressed the CDH trap and Shh-CB-msfGFP in PLMCs, induced synchronous release, fixed at 90 MPR using cytoneme-preserving Fidelis Fixative, and then imaged using Airyscan SR microscopy. As described above for un-trapped Shh-FL-msfGFP, we observed accumulation of Shh-CB-msfGFP along cytonemes and within large endosome-like structures (Figure 2G). We noted that these endosome-like structures are roughly ≥ 1μm in diameter and often contain smaller internal vesicles resembling the intralumenal vesicles of multivesicular bodies (Figure 2G, bottom right panel, Supplementary Figure 4C). To determine where and when these Shh+ endosome-like structures originate, we induced synchronous release of Shh-CB-msfGFP in PLMCs and used super-resolution spinning disc microscopy to image Shh trafficking live. Shh+ endosome-like structures (yellow arrowheads) begin to appear about 40 MPR, often near the Golgi apparatus, and they continued to accumulate over the next 20+ minutes (Figure 2H). These Shh+ endosome-like structures again are roughly ≥ 1μm in diameter, contain internal vesicles, and are observed moving throughout the cell body (Movie 4).

### Shh+ endosome-like structures are bona fide multivesicular bodies that can secrete exosomes into the extracellular space

Based on these observations, we hypothesized that these Shh+ endosome like structures that appear after release are multivesicular bodies (MVBs). MVBs are distinguished from other endosome-like structures by their enrichment of specific membrane proteins, such as the tetraspanin CD63, by their inclusion of smaller internal cargo-containing intralumenal vesicles, and by their endocytosis-dependent generation. (Figure 3A).^39,40^ To test whether Shh+ endosome-like structures share these properties, we first created a CDH compatible CD63 fusion protein by inserting the CB moiety in the first extracellular loop upstream of a mKate2 and then performed a CDH co-release of both Shh-CB-msfGFP and CD63-CB-mKate2 to examine their co-localization (Figure 3B). At 90 MPR, we observed strong colocalization of Shh and CD63, indicated by the high Pearson’s Coefficient, within individual endosomal membranes as well as small punctate vesicles clustered within the larger spherical structures, verifying the identity of these structures as MVBs (Figure 3B, inset, Figure 3C). We observed similar co-localization of several other Shh fusion proteins (Shh-FL-mScarlet or Shh-HaloTag) with CD63-pHluo (which labels MVBs at neutral pH) in PLMCs, indicating that colocalization Shh-CB-msfGFP with CD63-CB-mKate2 is not an artifact of the CB moiety or synchronous release (Supplemental Figure 3). To further test whether the formation of Shh + MVBs depend on endocytosis, we repeated the CDH release of Shh with and without expression of Rab5a(S34N), a dominant negative (DN) mutant GTPase that is known to inhibit endocytosis.^41,42^ In control releases, Shh-CB-msfGFP accumulates heavily in MVBs from 60-90 MPR (Figure 3D, left panels, inset). However, expression of Rab5a-DN(S34N) prevented Shh-CB-msfGFP endocytosis and formation of MVBs (Figure 3D, right panels) and resulted in high levels of Shh accumulation at the cell periphery Figure 3D, bottom right panel inset). From these data, we conclude that the Shh+ endosome-like structure observed upon release of trapped Shh are bona fide MVBs with canonical morphology, molecular identity, and endocytic origin.

**Figure 3:**
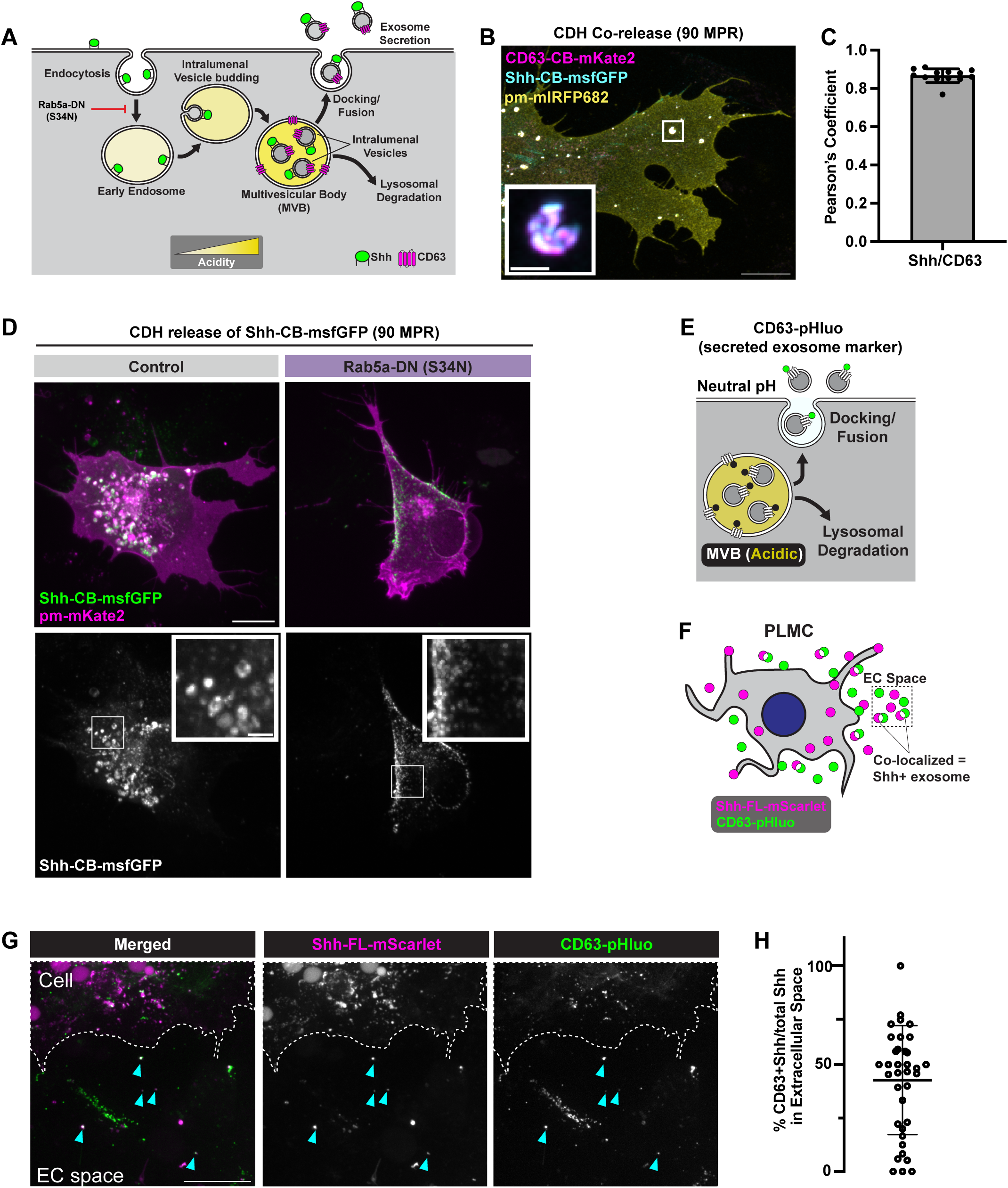
Endosome-like structures are bona-fide multivesicular bodies (MVBs) that fuse to plasma membrane to release Shh-bound exosomes. A) Diagram of exosome biogenesis within a cell, Shh (green) and CD63 (magenta). Molecules at plasma membrane are endocytosed and fuse with endosome. Endosome matures and begins budding inward creating the intralumenal vesicles of the MVBs. MVBs, labeled with the tetraspanin CD63, can either fuse to a lysosome and be degraded or fuse to the plasma membrane releasing its internal vesicles as exosomes. Note Rab5a-DN can inhibit the initial endocytosis step required for MVB generation. B) Airyscan SR Micrograph showing co-release of Shh-CB-sfGFP (cyan) and CD63-CB-mKate2 (magenta) in fixed PLMC co-labeled with pm-mIRFP682 (yellow) 90 MPR - Scale Bar = 10 μm. Inset highlights Shh+CD63+ MVB - Scale Bar = 1 μm. C) Graph showing high level of colocalization between Shh-CB-msfGFP and CD63-CB-mKate2 as depicted by high Pearson’s Correlation Coefficient (n = 13 cells, +/-SD). D) Micrographs CDH release of Shh-CB-msfGFP 90 MPR in control and PLMCs expressing Rab5a-DN(S34N) – Scale Bar = 10μm. Insets show Shh localizes to MVBs in control and to cell perimeter in Rab5a-DN(S34N). E) Diagram highlighting the secreted exosome marker CD63-pHluo (M153R). Note how the fluorescence of the pHluo is quenched in the lumen of the acidic MVB but will fluoresce on secreted exosomes exposed to the neutral pH of the extracellular (EC) space. F) Diagram showing experimental strategy to detect Shh colocalizing with CD63-pHluo. G) Micrograph of EC space outside neighboring a PLMC co-expressing Shh-FL-mScarlet and CD63-pHluo. Cyan arrowheads highlight EC colocalization - Scale bar = 10μm. H) Quantification of EC Shh-FL-mScarlet colocalization with CD63-pHluo (n = 36 cells/EC spaces, +/- SD).

MVBs can fuse to lysosomes, resulting in degradation, or can they fuse to the plasma membrane, resulting in the secretion of exosomes. To test whether Shh+ MVBs function to secrete Shh+ exosomes into the extracellular (EC) space, we first used superecliptic pHluorin (M153R) tagged CD63 (CD63-pHluo) as a proxy for secreted exosomes. pHluo fluoresces brightly in the EC space (neutral pH) but is quenched in the acidic lumen of MVBs in living cells (Figure 3E).^43,44^ We co-expressed Shh-FL-mScarlet and CD63-pHluo in PLMCs, imaged live, and then measured the extent of overlap between the mScarlet and pHluo dots in the local EC around PLMCs (Figure 3F). We found that over 40% of secreted Shh-FL-mScarlet overlapped with the secreted exosome marker CD63-pHluo in the EC space (blue boxes), indicating that exosome secretion makes a major contribution to the total pool of secreted Shh (Figure 3G, 3H). Overall, these data suggest that Shh is rapidly endocytosed from the plasma membrane after secretion, packaged into MVBs, and then secreted into the EC space bound to diffusible exosomes. Given the high degree of accumulation in MVBs and colocalization with secreted exosomes, we postulate that exosome packaging and diffusion is a key strategy for Shh dispersal in mesenchymal tissues.

### Exosome-bound Shh is transported along the external surface of cytonemes

While the above results suggest that much of secreted Shh is associated with diffusing exosomes, we also observe Shh moving along cytonemes, suggesting that these two modes of dispersal might somehow function together (Figure 1).^24^ One possibility, as previously proposed for Drosophila epithelia or cultured fibroblast cells^23,26^, is that Shh might travel within cytonemes, as Shh+ MVBs or exosomes, or in some other form. Through scanning electron microscopy, we observe that limb bud mesenchymal cell cytonemes are 100-200 nm in diameter while the size of MVBs have been reported to be 250-1000 nm,^45–47^ which makes the hypothesis that MVBs travel within cytonemes unlikely (Supplemental Figure 4). An alternative possibility is that secreted Shh+ exosomes bind to and travel along the external surfaces of cytonemes. Consistent with this possibility, in CDH release experiments, we observed large, spherical Shh structures (yellow arrowheads), presumably MVBs, travel to the base of a cytoneme, and then break apart into smaller Shh-containing puncta that undergo directed movements along the cytoneme towards its distal tip (yellow arrows) (Figure 4A, Movie 5). We hypothesized that these puncta represent newly secreted Shh+ exosomes that are transported along the external surface of the cytonemes. To further test this idea, we co-expressed Shh-FL-mScarlet, the secreted exosome marker CD63-pHluo, and pm-mIRFP682 to visualize Shh+ exosomes associated with cytonemes in living PLMCs. We found that roughly 40% of cytonemal Shh-FL-mScarlet colocalized with CD63-pHluo (white boxes) (Figure 4B-C). Additionally, we observe dynamic transport of exosome-bound Shh along PLMC cytonemes in both directions (Movie 6). Lastly, to confirm that exosome-associated Shh is traveling along the outsides of cytonemes, we performed immunofluorescence staining of extracellular (EC) Shh in non-permeabilized PLMCs (Figure 4D) and found that the vast majority of cytoneme-associated Shh is exposed to the extracellular space, implying that it is associated with the extracellular leaflet of cytoneme membranes (Figure 4E-F). This result is consistent with a previous study, which used split-GFP to detect association of Shh with the external surfaces of cytonemes in the limb bud.^24^ This result differs from the previously reported internal localization of Shh in fibroblast cytonemes, which suggests a potential cell-type specific mechanism of Shh transport along cytonemes.^26^ Altogether, we conclude that Shh+ exosomes not only diffuse in the EC space, but also, bind to and travel along the extracellular surface of cytonemes. This might act as a mechanism for directed dispersal or to increase the dispersal range of Shh+ exosomes.

**Figure 4:**
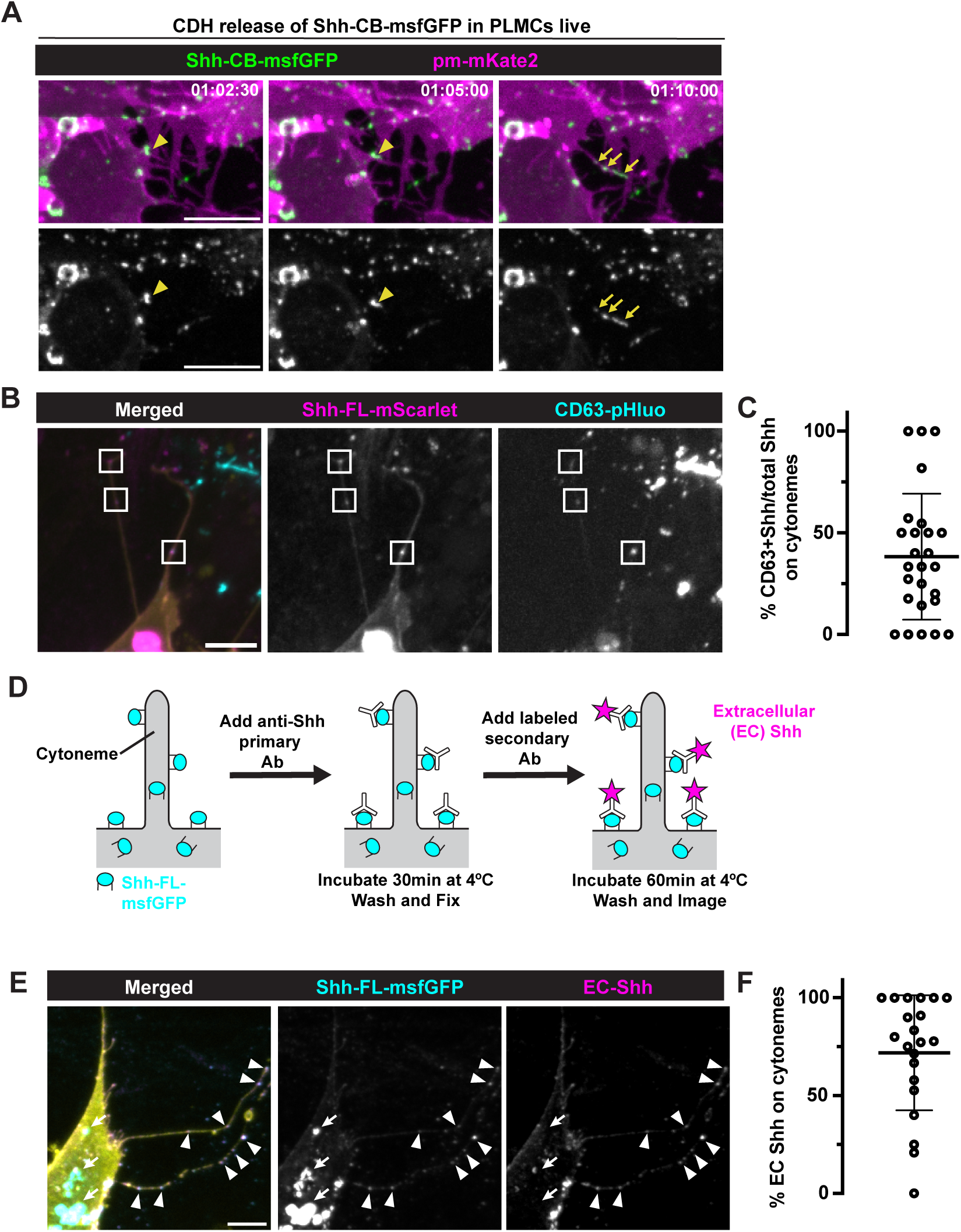
Shh-bound secreted exosomes can bind and travel along cytonemes. A) Timelapse of CDH release of Shh-CB-msfGFP in PLMC starting 1hour 2 min 30s post release. Note Shh+ MVB (yellow arrowhead) move to the base of a cytoneme and split apart into small Shh puncta (yellow arrows) along the cytoneme – Scale Bar = 10μm. B) PLMC co-expressing Shh-FL-mScarlet, CD63-pHluo, and pm-mIRFP682 (yellow) - Scale Bar = 5µm. On cytonemes, a fraction of Shh colocalizes with CD63-pHluo (white boxes). C) Quantification of colocalization (n = 26 cells, +/-SD). D) Diagram of extracellular (EC) Shh immunofluorescence staining protocol. E) EC Shh IF staining in PLMC - Scale Bar = 5µm. EC labeled Shh on cytonemes (white arrowheads). Note large MVB structures in cell body are unlabeled showing lack of internalization of antibody (white arrows). F) Quantification of colocalization between Shh-FL-msfGFP (Total) and EC-labeled Shh on cytonemes (n = 21 cells, +/-SD).

### Exosome secretion is required for 2D Shh signaling gradient formation

Our trafficking and localization data has suggested that exosome packaging and secretion may act as a key mechanism to disperse Shh and establish signaling gradients in mesenchymal tissues. Recent work, established a highly sensitive, reconstituted signaling gradient assay *in vitro* to probe Shh gradient formation.^15,48^ This reconstituted gradient assay is comprised of sparse “Sender” fibroblasts (Shh expressing) plated on a confluent layer of “Receiver” fibroblasts (Gli Binding Site fluorescent reporter) and, over time, a radial signaling gradient is formed around the Sender cell. We modified this assay so that the Sender cells are anterior limb bud derived PLMCs (no endogenous Shh expression) electroporated with a plasmid driving both Shh-FL-msfGFP and a palmitoylated fluorescent protein (Figure 5A). The tight correlation of expression between Shh-FL-msfGFP and membrane-FPs allowed us to use membrane FP intensity, as a proxy for Shh expression, to normalize gradient intensity in reproducible manner (Supplemental Figure 5). Overall, we reasoned this is a strong system to quantitatively test the role of exosome secretion in gradient formation in a 2D mesenchymal tissue.

**Figure 5:**
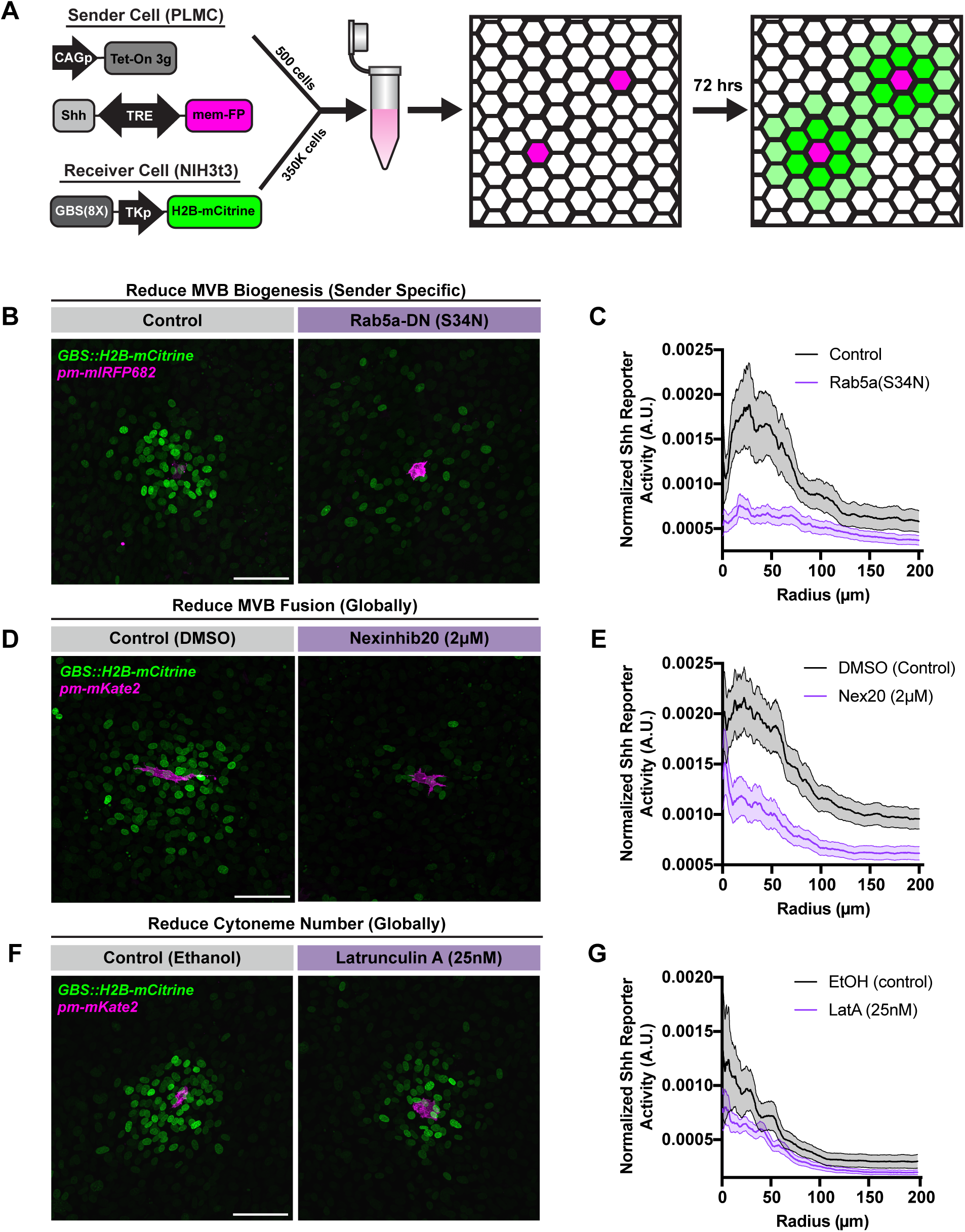
Exosome secretion is necessary for Shh dispersal and signal gradient formation. A) Diagram representing the experimental strategy of the reconstituted signaling gradient assay. Sender Cell PLMCs are electroporated with a plasmid encoding TetON3g and a plasmid that has a bidirectional Tet Response Element (TRE) promoter driving the expression of Shh-FL-msfGFP and a palmitoylated FP. B) Example gradients formed by control and Rab5a-DN(S34N) expressing PLMC sender cells. C) Graph plotting meta-gradients (i.e. average gradients) of control vs Rab5a-DN(S34N). Specifically, the mean normalized intensity +/- SEM of Shh Reporter Activity as a function of distance from center of the gradient is plotted (n = 17 control gradients, n =15 Rab5a-DN(S34N) gradients). D-E) Example gradient micrographs formed by DMSO and Nexinhib20 treated cells and accompanied quantification (n = 31 DMSO gradients, n = 30 Nex20 gradients). F-G) Example gradient micrographs formed by Ethanol and Latrunculin A treated cells and accompanied quantification (n = 24 Ethanol gradients, n = 27 Latrunculin A gradients). Scale Bars = 100μm.

To test the role of exosome secretion in gradient formation, we needed loss of function approaches that perturb exosome formation and/or secretion. We previously showed that expression of Rab5a-DN(S34N) greatly inhibited the biogenesis of Shh+ MVBs (Figure 3D). Using this tool, we inhibited MVB biogenesis in the PLMC sender cells and analyzed the resulting gradients. Here, we observe a dramatic reduction in Shh signaling gradient formation with local signaling (Shh Reporter Activity within 100 μm distance from gradient centroid) being almost eliminated completely (Figure 5B and 5C). However, we often observed slight activation of sparse Receiver cells many cell distances away from the centroid position suggesting Shh dispersal was still possible but not prevalent/strong enough to generate gradients.

Since blocking endocytosis is a relatively upstream manipulation of exosome biogenesis that could have off target effects, we aimed to block exosome secretion itself. Nexinhib20 (Nex20) is a small molecule inhibitor of the Rab27a GTPase that mediates MVB docking/fusion with the plasma membrane and has been reported to be a specific inhibitor of exosome secretion.^49,50^ As previously reported, we too found that Nex20 could reduce MVB fusion events in PLMCs (Supplemental Figure 6B-C, Movie 7). We repeated our gradient assays with either DMSO (control) or Nex20 incubation. Here, we found that Nex20 treatment greatly reduced the overall amplitude and intensity of signaling (Figure 5D and 5E). Altogether, these experiments demonstrate that exosome secretion is required for high-intensity shorter range (≤100μm) Shh dispersal that establish signaling gradients in 2D mesenchymal tissues.

Since cytoneme transport of Shh is seen in PLMCs, we also wanted to test the role of cytonemes in Shh dispersal and gradient formation. However, considering that PLMC cytonemes are very short in length, averaging between 2-10μm, we reasoned cytonemal dispersal alone could never establish a signaling gradient of this size (∼100 μm). So unsurprisingly, when we inhibit cytoneme number and length in all cells by incubation with a low concentration of Latrunculin A, an actin polymerization inhibitor that reduces filopodia formation (Supplemental Figure 6 D-F), we observe no alteration of Shh gradient formation (Figure 5F-G).^51–53^ This result validates previous reports showing that fibroblast cytonemes, of similar length to PLMC cytonemes, are dispensable in 2D gradient assays.^15^ So, despite seeing Shh accumulate and travel along cytonemes in PLMCs *in vitro*, the reduced lengths of these cytonemes also likely reduces their function in 2D gradient formation.

### Shh is likely dispersed via both exosome secretion and cytoneme transport in vivo

Considering that the limb bud mesenchyme is a disordered, 3D patterning field, it is likely that the dispersal mechanisms that govern 2D gradient formation might not be able to fully account for the added geometric and ECM complexity *in vivo*. It was previously reported that limb bud mesenchymal cytonemes could extend great lengths, up to 150μm, that can theoretically disperse Shh across the entirety of the 300 μm distance of the limb bud patterning field.^24^ To better compare the cytonemes of PLMCs and the *in vivo* limb bud mesenchymal cells, we performed *in ovo* electroporation of plasmids encoding pm-EGFP to visualize cytonemes in the developing limb bud mesenchyme (Figure 6A). Live imaging of limb bud mesenchymal cells *in vivo* shows a convoluted and interconnected network of cells projecting long cytonemes in 3 dimensions (Figure 6B). Like previous reports, we observe some stable cytonemes while others are dynamic both extending and retracting. After measuring over 300 cytonemes *in vivo,* we plotted and compared the lengths to the 2D PLMC cytonemes measured in Figure 1D. We can see that cytoneme length is far greater and the spread is much broader *in vivo* compared to *in vitro* PLMCs (Figure 6C). Overall, the greater number and length of cytonemes *in vivo* suggests a far more probable mechanism for long-range Shh dispersal and signal gradient formation.

**Figure 6:**
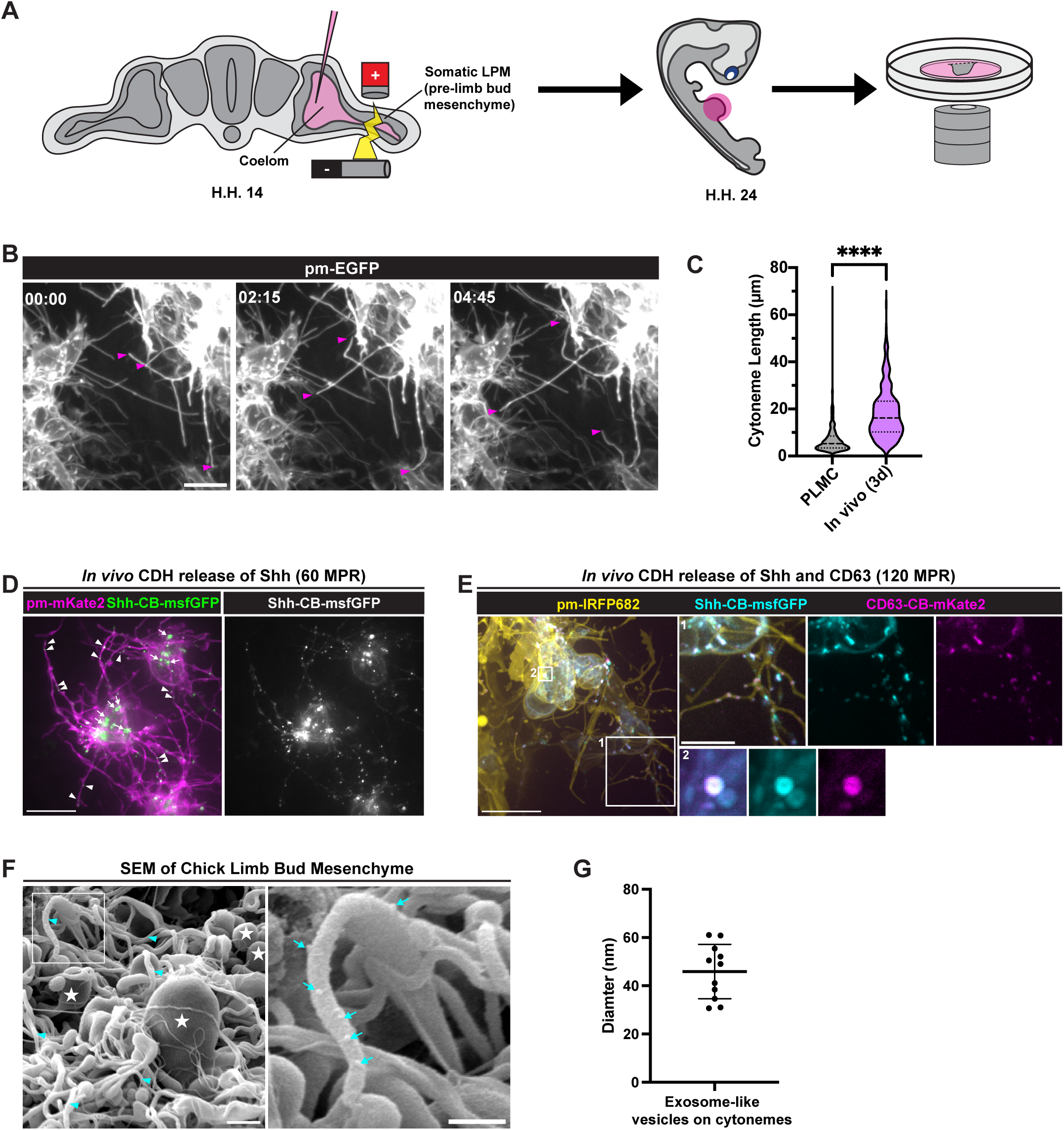
Shh-bound exosome secretion and cytonemal transport mechanisms are conserved in the limb bud mesenchyme *in vivo*. A) Diagram illustrating *in ovo* electroporation of the chick limb bud mesenchyme. B) Timelapse micrographs (minutes:seconds) of limb bud mesenchymal cells *in ovo* electroporated with pm-EGFP – Scale Bar = 10 μm. Note highly interconnected cytoneme networks between cells and the extension and retraction of long cytonemes (cytoneme tips labeled with magenta arrows). C) Quantification of 2D PLMC cytoneme lengths (From Figure 1D) compared to 3D cytoneme lengths *in vivo* (n = 490 PLMC cytonemes, n = 307 *in vivo* cytonemes, Mann-Whitney U test performed, p < 0.0001). D) Micrographs depicting CDH release of Shh-CB-msfGFP at 60 MPR *in vivo* – Scale Bar = 10 μm. Note Shh traffics along cytonemes (white arrowheads) and accumulates in MVB-like structures in cell body (white arrows). E) Micrographs depicting CDH co-release of Shh-CB-msfGFP and CD63-CB-mKate2 at 120 MPR *in vivo* - Scale Bar = 10μm. White box #1 shows magnified view of Shh+CD63+ exosomes traveling along cytonemes. White box #2 shows magnified view of Shh+CD63+ MVB in the cell soma. F) Scanning electron micrograph of intact limb bud mesenchyme in the ZPA region (HH 22-24). Note the globular cell bodies (white stars) and the complex network of cytonemes (cyan arrowheads) – Scale Bar = 1 μm. The white box highlighting one cytoneme in the left panel is magnified on the right panel. Note spherical exosome-like structures decorating the external surface of the cytoneme (cyan arrows) – Scale Bar = 500 nm.

Despite the considerable differences in cytoneme length, we sought to determine if Shh trafficking and overall distribution was conserved between PLMCs and limb bud mesenchymal cells *in vivo*. We first performed CDH releases of Shh-CB-msfGFP *in vivo* and imaged 60 MPR. Here, we observe Shh puncta decorate the shafts of these long cytonemes (Figure 6D, arrowheads). To note, more Shh-CB-msfGFP puncta localize to cytonemes *in vivo* compared to what we observe in PLMC cytonemes. This could imply a greater role of cytonemes in Shh transport *in vivo.* Additionally, like in PLMCs, we see Shh accumulate in MVB-like structures in the cell bodies of limb mesenchymal cells *in vivo* (Figure 6D, arrows). By performing co-releases of Shh and CD63, we show high levels of colocalization in these cell body MVBs Figure 6E, white box #2) and in exosomes traveling along cytonemes (Figure 6E, white box #1). To further validate this notion that exosomes travel along the external surface of cytonemes *in vivo*, we performed scanning electron microscopy on ectoderm peeled limb buds and observed small (30-60 nm), spherical, vesicles along a subset of cytonemes in the ZPA region (Figure 6F-G). Considering shared size and similar morphology to scanning electron micrographs of purified exosomes, we hypothesize that these vesicles are bona fide exosomes traveling along cytonemes which validates our previous light microscopy data. Altogether, these experiments verify that the dispersal mechanisms we observe in PLMCs are conserved *in vivo*.

Lastly, we pose a model in which there are two interconnected modes of exosome mediated Shh dispersal in the limb bud mesenchyme. First, secretion of Shh-bound exosomes into the EC space can function in high intensity signaling at a shorter range from the Shh-producing cells of the ZPA. Second, Shh-bound exosome transport along cytonemes can function in longer-range or directed, lower intensity signaling.

## DISCUSSION

The mechanisms that govern morphogen dispersal and gradient formation are central to understanding developmental patterning but remain poorly understood. Our study establishes a fundamental role for exosome secretion in mediating Shh dispersal and patterning in the limb bud mesenchyme. Using a combination of novel trafficking assays, synthetic tissue reconstitution, and quantitative *in vitro* and *in vivo* microscopy, we found that exosome secretion serves as a primary mechanism for Shh packaging, extracellular dispersal, and for the establishment of tissue-level signal gradients. While secretion and diffusion of Shh-bound exosomes is sufficient to form short-range (∼100 µm) signaling gradients *in vitro*, we also observe that secreted Shh-bound exosomes can bind to and move along cytonemes both *in vitro* and *in vivo*, suggesting a potential basis for longer-range and/or directed transport of Shh *in vivo*. Thus, the combination of exosome-mediated secretion with different modes of exosome transport could provide a robust and versatile way to sculpt gradients of Shh signaling activity across different developing tissues with different architectures, geometries, and across different spatial and temporal scales.

### CDH Synchronous Release System provides a powerful tool to visualize morphogen trafficking in embryonic tissues

An ability to visualize secretory traffic is key to determining when and how morphogens are packaged for secretion and dispersal. Previous trap and release methods used to visualize trafficking are based on the use of exogenous biotin/streptavidin to trigger release of trapped cargo. However, the presence of endogenous biotin has limited the use of these methods in vertebrate embryonic tissues, which contain high levels of endogenous biotin. To overcome this limitation, we developed an alternative strategy, based on chemically disruptable heterodimers (CDH): using rationally designed protein off switches comprised of Bcl-xL receptors and synthetic binding moieties to construct the trap, and small molecule inhibitors to induce synchronous release. We confirmed that this approach allows us to trap and release Shh in both primary cultures and embryonic vertebrate tissues *in vivo* with limited leak (Supplemental Figure 2), and we used the approach here to visualize and quantify Shh flux through the secretory pathway, revealing a key role for multivesicular body packaging to prime the cell for Shh+ exosome secretion (Figure 2, 3). More generally, the CDH synchronous release system offers a powerful and versatile approach with unparalleled control to study morphogen trafficking and dispersal dynamics in native, tissue environments *in vivo*.

### Exosome secretion is critical for establishing Shh signaling gradients in mesenchymal tissue

Shh has been shown to associate with MVBs and exosomes in cultured immortalized cells and epithelial tissues ^20–23^, and inhibiting endocytosis in Hh-producing cells can reduce the spatial extent of signaling.^20–23,54^ However, the contributions of exosome-based secretion to shaping Hh signaling gradients in all tissues, and especially in mesenchymal tissues remain poorly understood. Here we have shown that accumulation of Shh in multivesicular bodies, followed by exosome release, is a predominant pathway for Shh secretion in limb bud mesenchymal cells. We find that a substantial fraction of newly synthesized Shh is rapidly trafficked into MVBs after synthesis in PLMCs and *in vivo* (Figure 2G-H, 3B, 6D-E), and over 40% of Shh in the EC space co-localizes with CD63-pHluo, a marker for secreted exosomes (Figure 3H). Using a synthetic tissue assay in which Shh reporter cells read out gradients of Shh signaling activity, we have shown that Shh released from individual PLMCs can form high intensity and short-range signaling gradients (∼100 μm) *in vitro* (Figure 5). These gradients are sharply attenuated by inhibiting endocytosis, or by inhibiting MVB fusion and exosome release, and they are insensitive to inhibition of cytonemes, confirming that they are formed by secretion and extracellular diffusion of Shh-bound exosomes. Although limb bud mesenchymal cells express other extracellular transporters of Shh, such as Scube1 and Scube3,^55,56^ their presence is not sufficient to sustain Shh signaling gradients in our synthetic tissue assay when exosome-based secretion is inhibited. Thus altogether, our results suggest that Shh, carried by secreted exosomes, plays the dominant role in shaping Shh signaling gradients *in vitro*, although additional carriers, such as Scube proteins, or cytoneme-based transport, may contribute to shaping Shh gradients in vivo.

### Transport along cytonemes provides an alternative mode of exosome-mediated Shh dispersal

Previous studies showed that Hh co-localizes with exovesicle/exosome markers on cytonemes in immortalized fibroblasts and in Drosophila wing disc epithelia.^23,26^ However, the origins of these Hh+ exovesicles, and whether they travel on or within cytonemes, remains unclear. Here, through synchronized release, live imaging of CD63-pHluo, and IF staining of extracellular Shh, we showed that a significant fraction of secreted Shh+ exosomes can bind to and travel along the external surface of cytonemes in PLMCs. This suggests that the same secreted Shh-containing exosomes can be dispersed by diffusion through extracellular space or by directed transport along cytonemes. Our observations of co-released Shh and CD63 within the intact limb bud mesenchyme reveal that Shh+ exosomes can also bind to and move along cytonemes *in vivo*, suggesting a role for cytonemal transport of Shh+ exosomes during normal development. While inhibiting cytonemes has no effect on Shh signaling gradients in our synthetic tissue assays, the cytonemes formed by PLMCs *in vitro* are much shorter than the spatial extent of Shh signaling (∼100 µm) observed in these experiments; thus, the contribution of exosome diffusion is likely to dominate in these *in vitro* assays. In contrast, limb bud mesenchymal cells project long, dynamic cytonemes *in vivo,* that can span the entire 300 μm limb bud patterning field, providing far longer tracks for exosomes to travel along (Figure 6).^24^ Furthermore, exosome diffusion may be restricted in the more complex 3D extracellular environment of the limb bud *in vivo*. Thus, transport of Shh-containing exosomes along cytonemes *in vivo* may help to extend Shh signaling gradients beyond the range accessible by extracellular diffusion.

Based on these observations, we propose that local diffusion and cytoneme-based transport of Shh-containing exosomes work together to shape spatial gradients of Shh signaling in the limb bud. Studies in chick wing buds demonstrated that Shh concentration and duration control digit number and identity, consistent with classical morphogen models.^57,58^ Local diffusion of Shh+ exosomes away from source cells shapes short-range, high intensity signaling gradients that can explain the specification of more local posterior fates, while transient binding and directed transport of Shh+ exosomes along cytonemes could provide longer-range low intensity Shh signaling that specifies anterior digit fate. Furthermore, recent work in mouse models have identified two phases of Shh signaling: 1) a short burst of more local ZPA signaling that patterns the most posterior digits 2) a secondary phase of longer range signaling that promotes expansion, survival, and indirectly specifies more anterior digits.^59,60^ In this context, rapid exosome diffusion away from source cells could provide the transient burst that is necessary and sufficient for posterior digit specification, while transport of Shh+ exosomes along cytonemes could supply longer-range and delayed Shh signaling required for the secondary wave of expansion and cell survival.

### Exosome secretion may unify multiple modes of Shh dispersal

Our work identifies exosomes as a major carrier of secreted Shh and highlights how the combination of diffusive and directed transport of exosomes could sculpt Shh gradients in different ways^23,26^ However, it is likely that many other factors contribute to shaping the dispersal and bioactivity of exosome-associated Shh. In other contexts, exosomes have been shown to carry Dispatched,^23^ which may act to present Hh to other extracellular transporters like Scube or HDL in the extracellular space. Thus, it will be interesting in future work to define the composition of Shh signaling machinery on limb bud mesenchymal cell exosomes, and how these contribute to shaping the spatiotemporal dynamics of Shh signaling, or to endowing individual exosomes with different signaling capacities, allowing them to play different roles in mitogenic vs morphogenic signaling, as has been proposed in other contexts.^20,54^

Speaking more broadly, studies of Hh signaling have largely focused on characterizing the individual contributions of different factors to shaping spatial gradients of Hh signaling activity in different developmental, physiological and organismal contexts. However, a growing body of work, including ours, supports the broader view that multiple factors work together to control the secretion, dispersal and bioactivity of Hh and that their relative contributions are tuned differently in different contexts to shape the spatiotemporal dynamics of Hh signaling. Our work raises the exciting possibility that secreted exosomes could play a central role for integrating these diverse inputs into coherent Hh signaling programs. Looking beyond tissue patterning, intercellular signaling is also a major driver of disease progression. Many developmental signaling programs are reactivated in diseases such as cancer.^61^ Previous work has highlighted the importance of exosome-mediated^62,63^ and cytoneme-mediated^64,65^ signaling among tumor cells in patterning their activities and in the broader tumor microenvironment. Thus, better understanding of how these and other regulators of Hh dispersal and bioactivity work together to pattern surrounding tissues should yield insight into both development and disease.

### STAR★Methods

**Key resources table**

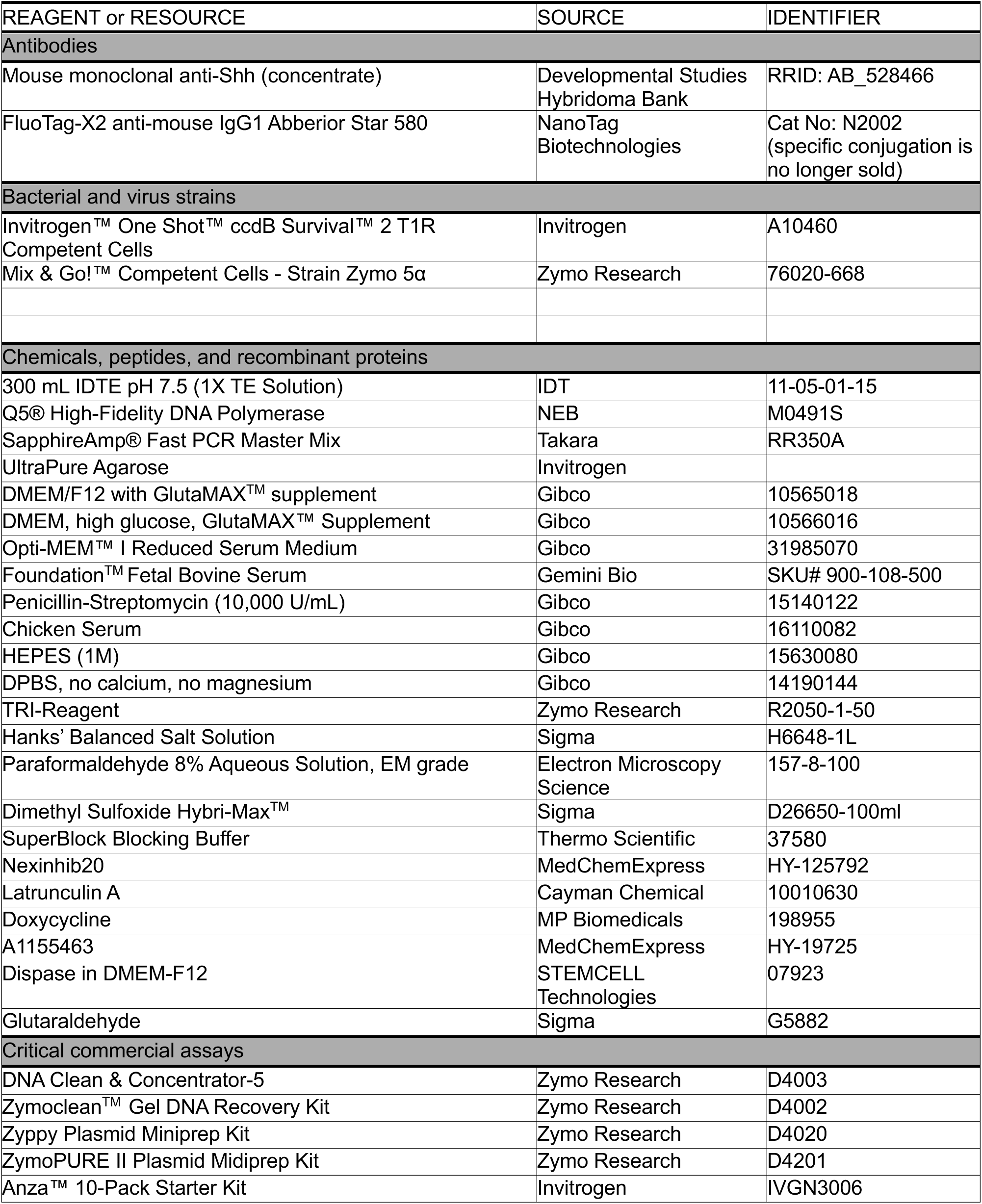
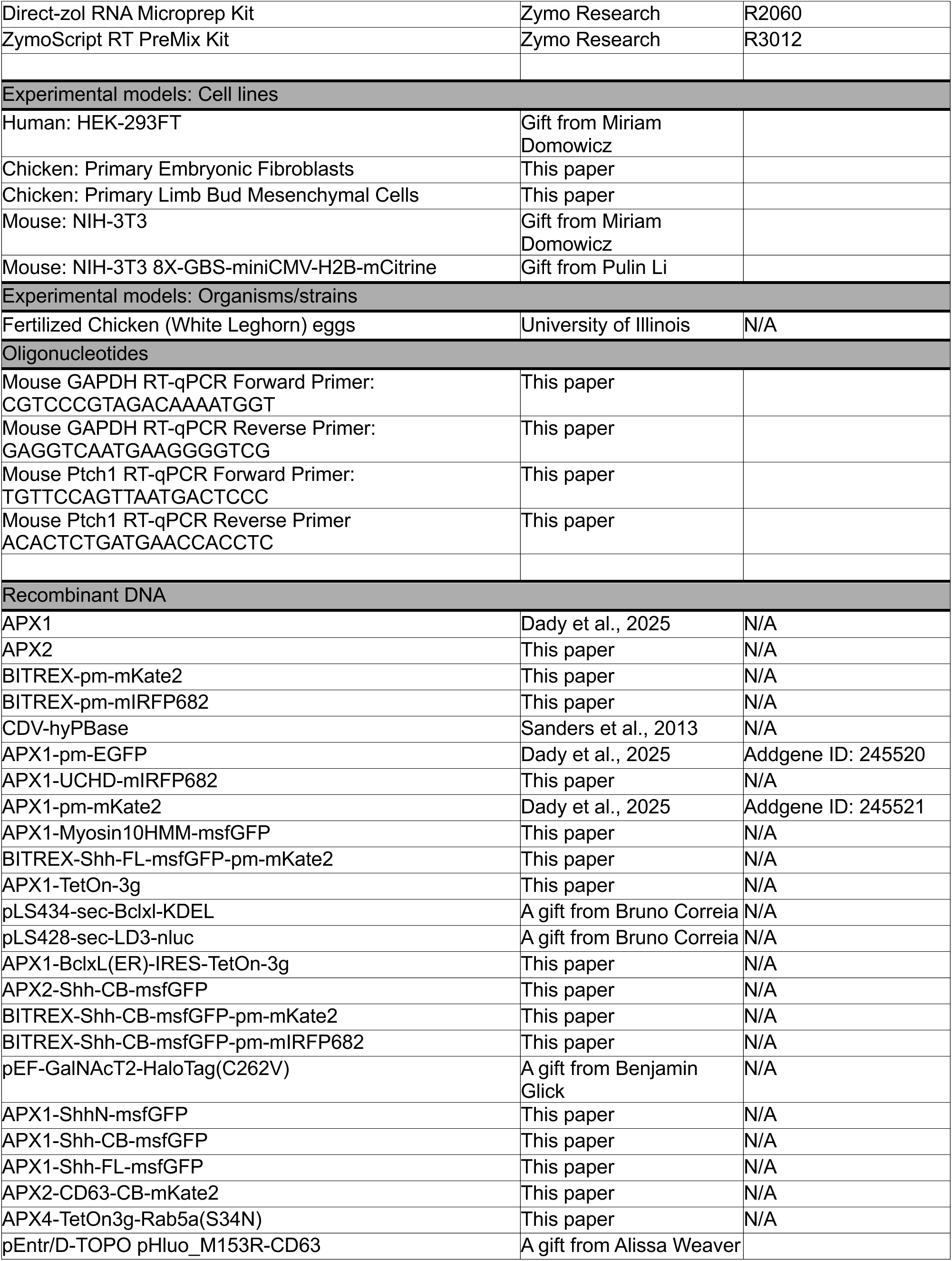
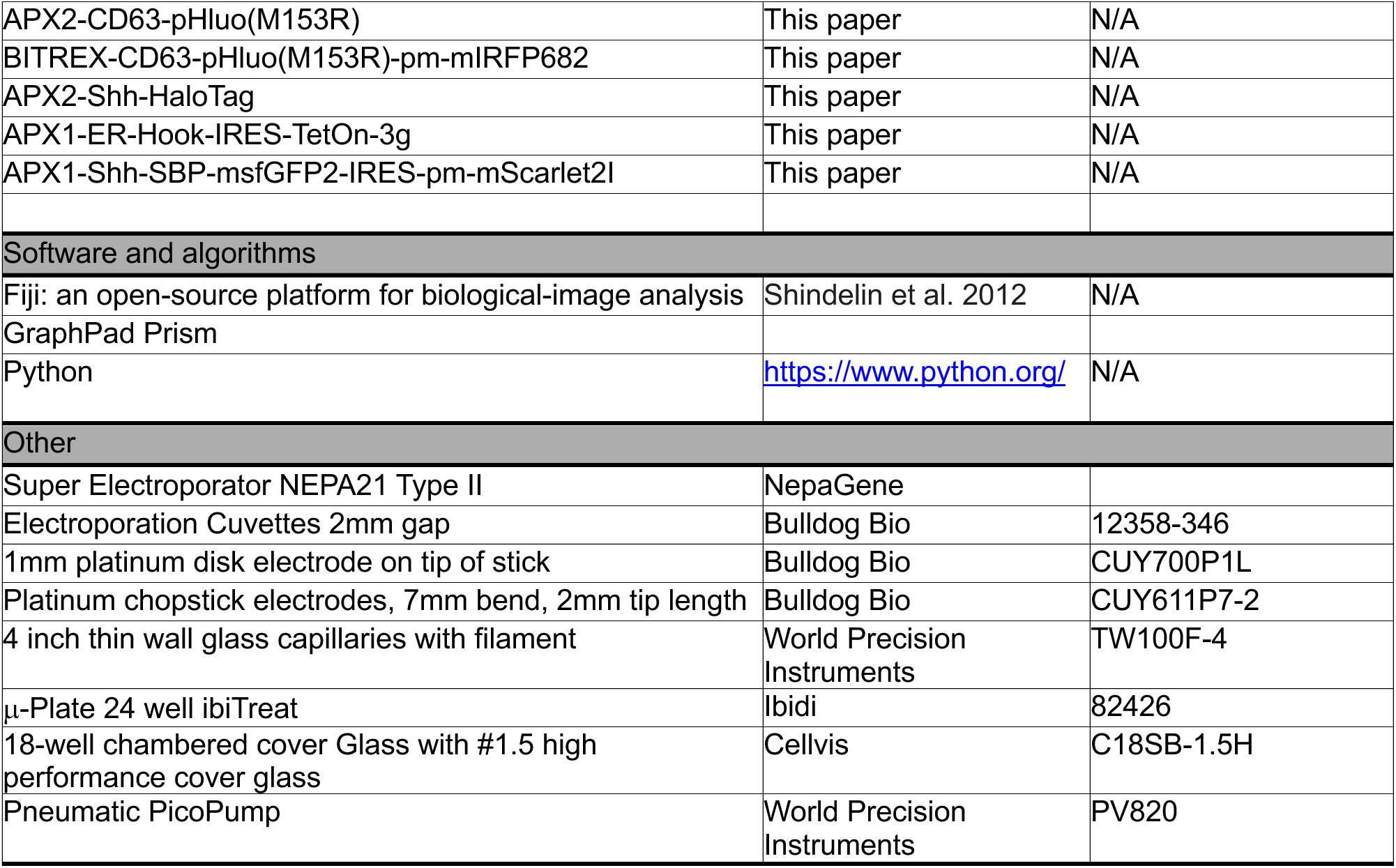

## EXPERIMENTAL MODEL

### Chick Embryos

We purchased fertilized White Leghorn chicken eggs (G. gallus) from the University of Illinois, Urbana-Champaign and stored them at 15°C until incubation. We incubated eggs in a non-rotary incubator at 37.5°C with approximately 50-60%-humidity until they reached the desired stage according to Hamburger and Hamilton (HH).^66^

### Generation of Primary Limb bud Mesenchymal Cells (PLMCs)

To generate PLMCs, we adapted a micromass culture approach.^24,67,68^ First, we incubated chicken eggs until they reached HH 22 (∼E3.5). We transferred embryos to room temperature Hanks’ Balanced Salt Solution (HBSS, Sigma) supplemented with penicillin/streptomycin (Gibco) and dissected out the limb buds using sterile forceps and Vannas scissors. We transferred all the limb buds to a microcentrifuge tube containing 1mL of Tryp-LE (Gibco) and incubated the tube for 20 minutes at 37°C. We then replaced Tryp-LE with DMEM-F12 (Gibco) supplemented with Pen/Strep, 10% FBS (Gemini), and 25 mM HEPES (Gibco). We used repeated passage through a 21G needle to fully dissociate limb buds into single cells. We then passed the dissociated cells through a 40 μm cell strainer, spun them down for 3 minutes at 200g, resuspended in fresh DMEM-F12 and then seeded 1 x 10^5^ PLMCs into each well of a glass-bottom 18-well chamber slide (CellVis). We incubated these cells at 37°C with 5% CO_2_. For some experiments, we introduced plasmids by electroporation before seeding (see *In Vitro* Electroporation).

### Generation of Chicken Embryonic Fibroblast (CEF)

We prepared cultures of Chick Embryo Fibroblasts (CEFs) similar to previous reports.^69^ All operations were performed under sterile conditions. Briefly, we used forceps to extract chicken embryos at embryonic day 10 (HH 36), removed the head, limbs and viscera, and used a razor blade to mince 4 - 6 embryos in a glass petri dish, before adding10 mL of Tryp-LE (Gibco). We used a wide-mouthed pipet to transfer the minced embryos in Tryp-LE to a sterile 250 ml flask, stirred gently for 15 minutes, allowed them to settle for 5 minutes, and then transferred the supernatant to a 50 mL conical tube. After vigorously pipetting the supernatant to dissociate any remaining tissue, we added an equal volume of 100% FBS and mixed by inversion to inactivate Tryp-LE. We allowed the solution to settle for 5 minutes, we transferred the supernatant to a fresh 50 mL conical tube, centrifuged for 5 minutes at ≤ 200g, removed the supernatant, and then resuspended in 10 mL of 100% FBS. After a second round of centrifugation (5 minutes at ≤ 200g) and removal of supernatant, we resuspended the pelleted cells in 5 mL of CEF Medium (10% FBS, 2% heat inactivated chicken serum (Gibco), in DMEM (Gibco; high glucose, GlutaMax, sodium pyruvate)). We then plated cells at various concentrations (between 10^5^ and 10^6^ cells) in 10 mL CEF Medium on 10 cm plates and incubated them at 37°C with 5% CO2. We passaged cells every 2-3 days and selected the healthiest cultures for further use and subsequent freezing.

### Cell lines

We routinely cultured HEK-293FT and NIH-3T3 cells were routinely cultured in DMEM high-glucose (Gibco) supplemented with Glutamax and 10% FBS (Gemini Bio) at 37°C and 5% CO _2_ according to standard procedures.

## METHOD DETAILS

### Plasmids

#### Plasmid vectors for constitutive and inducible expression

We used the piggyBac transposition system to genetically integrate DNA constructs for extended expression in cell culture and during chicken embryogenesis. We drove expression of the hyperactive piggyBac transposase using the CDV-hyPBase plasmid in all transfection/electroporation-based experiments.^70^

For stable, constitutive expression of various constructs, we used the APX1 backbone.^71^ This allowed for convenient Gateway mediated generation of numerous piggyBac compatible expression vectors with a strong constitutively active promoter. We created the APX2 destination vector by replacing the CAG promoter with a Tetracycline Response Element (TRE) to drive doxycycline inducible expression of genetic constructs. For bidirectional doxycycline inducible expression, we used the BITREX plasmid backbone which used a bidirectional TRE to drive dual expression of (a) either palmitoylated mKate2 or mIRFP682 and (b) an RFA Gateway cassette for Gateway LR Clonase II mediated recombination.

For TetOn-3g expression, we created two different destination vectors to enable expression of two genetic constructs from the same plasmid. First, we inserted an IRES2 element into the APX1 vector, downstream of the gateway cassette, to bicistronically co-express TetOn-3g with other constructs of interest. Second, we created a bidirectional backbone, APX4, that uses a Ubiquitin C (UBC) promoter to constitutively express TetOn-3g while a CAG promoter drives expression of a RFA Gateway cassette for convenient LR Clonase II mediated recombination from Gateway Entry Vectors.

#### Plasmids encoding the Chemically Disruptable Heterodimer (CDH) Synchronous Release System

To create the endoplasmic reticulum (ER) localized Bcl-xL trap portion of the CDH synchronous release system, we first cloned the sequence encoding human Bcl-xL sequence (gift from Bruno Correia). Then we used overlap extension PCR, to insert the first 38 amino acids (including the signal peptide) from the human STIM1L gene at the 5’ end of Bcl-xL and the nucleotides encoding a KDEL amino acid sequence at the 3’ end of Bcl-xL.

To create Shh-CB-msfGFP, we used overlap extension PCR to generate a construct containing, in order: the N-terminal domain of murine Shh, a flexible linker, the LD3 sequence (gift from Bruno Correia) with D138A point mutation (now deemed CDH Binder or CB), a mini linker, msfGFP, and a mini linker followed by the C-terminal domain of murine Shh to ensure proper autocatalytic processing and cholesterol modification.

To create CD63-CB-mKate2, we inserted both the CB and mKate2 moieties into the first extracellular loop of human CD63 into a previously published insertion site.^72^

Key vectors created and used for this work are deposited in Addgene, Inc.

### *In Vitro* Electroporation

For *in vitro* electroporation, we suspended 1 x10^6^ cells (PLMCs or CEFs) in 2mm gap cuvettes (Bull Dog Bio) containing 10 μg of plasmid DNA in 100 µL of a modified Sucrose Electroporation Buffer (68 mM Sucrose, 250 μM MgCl_2_, Opti-mem)^24,68^. We placed the cuvettes on ice for 5 minutes and then electroporated using a Nepa21 type II electroporator using a square wave pulse train of 5 poring pulses (275 V, 0.1 msec Pulse Length, 100 msec Pulse Interval, 10% Decay Rate, + Polarity) and 5 transfer pulses (20 V, 50 msec Pulse Length, 50 msec Pulse Interval, 40% Decay Rate, +/- Polarity). We then allowed incubated cells to recover on ice for 5 minutes before seeding onto glass bottom 18 well chamber slides (Cellvis) or tissue culture treated dishes (Ibidi).

### Chicken Embryo Microinjection and *In Ovo* Electroporation

We incubated fertilized chicken eggs at 37.5 °C for 50-52 hours until HH 13-14. To access embryos, we opened windows in the eggshell, added HBSS, and then gently dissected the vitelline membrane above the limb field. We used a Leica Stereoscope equipped with a 470/40 nm bandwidth emission filter to provide the necessary contrast against the background yolk for injection. We prepared plasmids for injection by dilution to 3-5 μg/μL in endotoxin free water, supplemented with 0.1M phenol red to enable visualization of injections. We loaded the plasmid solution into 1mm diameter glass capillary needles and then microinjected, using a Pneumatic PicoPump (WPI), into the embryonic coelom of the forelimb field. To electroporate plasmids into the somatic lateral plate mesoderm, we positioned a 2mm tip platinum chopstick negative electrode (Bulldog Bio) ∼1 mm below the embryo’s forelimb field and a 1mm platinum disk on a stick positive electrode (Bulldog Bio) 1-2 mm above the embryo’s forelimb field. We then used a Nepa21 electroporator (Nepagene) to deliver a square wave pulse train of one 8V pulse (50 msec pulse length) followed by two 6V pulses (50msec pulse length, 1 sec interval), then removed the electrodes and pipetted 500 μL of HBSS onto the embryo. Following microinjection or electroporation, we taped the egg window using packaging tape and transferred the egg to a 37.5°C incubator until embryos reached HH 21-23. We then dissected right forelimb buds and prepared them for either live or fixed imaging as described below.

To perform live imaging, we transferred dissected right forelimb buds into a glass bottom 35mm dish (Cellvis) containing DMEM-F12 supplemented with 10% FBS and 20mM HEPES. To immobilize limb buds for imaging, we used a ring of Vaseline to sandwich them between a 12 mm circular cover slip and the glass bottom coverslip of the 35mm dish. We performed all live imaging in 37°C heat chambers.

### Stable Integration and Doxycycline Regulated Gene Expression

#### Stable Integration of DNA constructs

For all microinjection/electroporation experiments, we added the piggyBac transposase plasmid (CDV-hyPBase) to the plasmid solution as a 0.1-0.5 molar fraction of the expression plasmids to mediate stable genomic integration and high expression over time.

#### In Vitro Gene Expression

For all experiments involving doxycycline induced expression of TRE-driven constructs, we introduced doxycycline (1-2 μg/mL) to the culture medium 24 hours post-electroporation/transfection.

#### In Ovo Gene Expression

For all experiments involving inducible TetOn-3g gene expression, we removed windowed eggs from incubation 24 hours after electroporation and injected 500 μL of doxycycline solution in HBSS (200 μg/mL) beneath the vasculature using a 27-gauge needle. We then incubated eggs for an additional day before further analysis.

### Labelling Cells with HaloTag Ligands

Cells expressing HaloTag fusion proteins were incubated in their respective their cell media supplemented with 500 nM Janelia Fluor JFX650 overnight. We then washed cells with fresh media prior to live or fixed imaging.

### Fixation

#### Fixation of cultured cells

To preserve cytonemes, we used a fixative cocktail, Fidelis Fix, containing 0.1M PHEM, 2% paraformaldehyde, 0.2% glutaraldehyde, and ddH20. We rinsed cells once with ice cold 0.1M PHEM buffer (Electron Microscopy Sciences), incubated them in Fidelis Fix for 15 minutes on ice, rinsed them once with ice cold PHEM buffer and then washed twice for 5 minutes each with PBS before further analysis.

To preserve cytonemes in dissected limb bud tissue, we pre-incubated limb buds in ice cold 0.1M PHEM buffer supplemented with 60 mM sucrose (S-PHEM) for 5 minutes. We then incubated limb buds in Fidelis Fix supplemented with 120 mM sucrose and on ice for 1 hour, washed once with S-PHEM buffer for 10 minutes, then washed three times with PBS for 15 minutes each before further analysis.

For all other experiments, we rinsed cells briefly with ice cold DPBS without calcium or magnesium (Gibco) and then fixed them using 4% paraformaldehyde (Electron Microscopy Sciences) in PBS on ice for 15 minutes, followed by three washes with PBS for 5 minutes each.

### Bioactivity Assay

#### Treating NIH 3T3 cells with conditioned media

We seeded 1 x 10^5^ HEK293FT cells into each well of a 12 well tissue culture dish containing DMEM supplemented with 10% FBS (DMEM10), cultured them for 24 hours at 37¼C, and then transfected each well with either APX1-pm-EGFP, APX1-Shh-N-msfGFP, APX1-Shh-FL-msfGFP, or APX1-Shh-CB-msfGFP using the DreamFect Gold kit (OZ Biosciences) according to the manufacturer’s protocol. After culturing for an additional 24 hours, we dissociated transfected cells using Tryp-LE as described above, centrifuged for 3 minutes at 200g, resuspended with 5 mL of DMEM supplemented 2% FBS (DMEM2), and transferred the dissociated cells to a 60mm tissue culture dish. After an additional 72 hours of culture at 37¼C, we collected the conditioned media from each dish, centrifuged for 5 minutes at 200g, and then filtered through a 0.22 μm filter. We added 2ml of freshly collected conditioned media, combined with 2 mL of new DMEM10, to NIH-3T3 cells grown to 75% confluency on 6 well dishes. We then cultured the cells in conditioned media for 40 hours before RT-qPCR analysis.

*RNA isolation, cDNA synthesis, and quantitative PCR:*

After 40 hours of culture in conditioned media, we performed RNA extraction using the Direct-zol RNA Microprep kit, followed by reverse transcription, using the ZymoScript RT PreMix Kit, to generate 50 ng/μL cDNA. We performed qPCR using PowerUp SYBER Green Master mix on a QuantStudio3 Real-Time PCR system (Applied Biosystems) for 40 cycles to amplify and measure murine *PTCH1* and *GAPDH* mRNA transcripts. We used the delta delta Ct (ddCt) method to quantify changes in relative gene expression induced by conditioned medium.^73^ Graphs display 2^-ddCt^ fold change and we computed statistical significance using one-way ANOVA with Tukey’s multiple comparisons test on dCt values.

### Confocal Microscopy

We acquired super-resolution confocal images using either a 3i Marianas Super-Resolution by Optical Reassignment (SoRa) Spinning Disk confocal, or a Zeiss LSM 980 Laser-scanning confocal equipped with airyscan2. For SoRa acquisitions, we used 100x/1.3 NA, Plan NeoFluor oil, 63x/1.4NA Plan Apochromat oil, and 40x /1.3NA Plan NeoFluor oil lenses (Zeiss), and recorded images on a Prime 95B Back Illuminated Scientific CMOS camera. For For LSM 980 acquisitions, we used a 63x /1.4NA Plan Apochromat oil lens (Zeiss) and acquired images in airyscan2 mode. Please see individual figure legends for further details about image acquisition parameters.

We acquired non-super-resolution confocal images using a Nikon Ti-E inverted microscope equipped with a Yokogawa CSU-X1 spinning disk scan head, and a Ti-ND6-PFS Perfect Focus Unit. A laser merge module (Spectral Applied Research) controlled tunable delivery of 481-nm and 561-nm laser excitation from 50 mW solid-state lasers (Coherent Technology). All image acquisition was controlled by Metamorph software. We collected images using a 60X Nikon Plan Apo 1.2NA water immersion objective onto an Andor iXon3 897 EMCCD camera.

### Total Internal Reflection Fluorescence (TIRF) Microscopy

We performed TIRF microscopy on a Nikon Ti2-E inverted microscope equipped with a LunF XL laser combiner housing solid state 488nm, 561nm and 640nm lasers and feeding three separate TIRF illumination arms, a Ti2 N-ND-P Perfect Focus unit, and a CFI APO 100X 1.49NA TIRF (Nikon) objective. We acquired images on an Andor IXON Life 897 EM-CCD camera to achieve a pixel size of ∼160 nm. Image acquisition was controlled by Nikon Elements software.

### Scanning Electron Microscopy (SEM) of Chicken Limb Buds

We dissected limb buds from HH 22-24 chicken embryos, incubated the limb buds in Dispase/DMEM-F12 solution for 60 minutes at 37 °C, and used forceps to gently remove the outer sheath of ectoderm. We then immediately fixed the limb bud mesenchyme using cytoneme preserving Fidelis Fixative with sucrose. To ensure tissue rigidity for EM processing, we then double fixed the limb bud mesenchyme in 2% paraformaldehyde and 2.5% glutaraldehyde in PHEM buffer. We then dehydrated the tissue in a series of ethanol solutions and transferred the tissue to the UIC EM staff. They fully dehydrated the limb buds using HMDS, mounted the limb buds onto an SEM pin with conductive tape, and sputter coated the tissue with 10 nm of platinum. We then imaged the limb bud mesenchyme at 5.0 kV using the Secondary Electron Detector on a JEOLJSM-IT500HR SEM. This work made use of instruments in the Electron Microscopy Core of UIC’s Research Resources Center.

### Image Analysis and Visualization

We used ImageJ/FIJI software to process all image data for analysis and display. We used standard maximum intensity projections to produce single projected views from confocal Z-stacks, cropped the images to highlight features of interest, and adjusted brightness and contrast, before saving final images in RGB format.

### Kinetic Analysis of Synchronously Released Shh

For synchronous release experiments, we co-electroporated CEFs with APX1-BclxL(ER)-IRES-TetOn3g, APX2-Shh-CB-msfGFP, and pEF-GalNAcT2-HaloTag(C262V). 48 hours after electroporation, we added media supplemented with 20μM A1155463 (A1155) to the cell culture dishes to induce synchronous release of Shh. We then performed timelapse imaging using the Marianas SoRa spinning disk confocal, and a 63x objective. Starting 5 minutes post-release, we acquired Z-stacks through entire cells (∼10 μm at 1 µm intervals in Z) every 3 minutes for one hour.

To analyze the kinetics of Shh release, we used ImageJ/FIJI to construct maximum intensity projections from the individual Z-stacks. We used a custom written FIJI macro (provided in supplement) to create ROIs of the Golgi apparatus at every time point and measure the integrated density of Shh-CB-msfGFP in each ROI. For comparison across cells, we performed min max normalization and plotted the normalized integrated density as a function of time.

### Cytoneme Quantifications

To quantify cytoneme numbers and lengths in PLMCs expressing palmitoylated fluorescent proteins (pm-EGFP, pm-mKate2), we used the Marianas SoRA to acquire Z-stacks through entire cells (∼10 μm at 0.5 µm intervals in Z). We produced maximum intensity projections and then measured cytoneme number and length using FIJI. To quantify cytoneme numbers and lengths for limb bud cells expressing palmitoylated fluorescent proteins (pm-EGFP, pm-mKate2) *in vivo*, we acquired Z-stack images of fields of cells then used the Neuroanatomy SNT plugin to measure and track cytonemes in 3D.^74^ In both cases, we considered all filopodia above 2 microns in length to be cytonemes.

To quantify cytoneme diameter in SEM images of the limb bud mesenchyme *in vivo*, we measured cytoneme width in three locations per cytoneme using FIJI and then averaged the measurements to create a composite diameter measurement.

### Analysis of Colocalization Following Co-Release of Shh and CD63

To measure co-localization of co-released Shh and CD63, we electroporated PLMCs with AXP1-Bclxl(ER)-IRES-TetOn3g, BITREX-Shh-CB-msfGFP-pm-mIRFP682, and APX2-CD63-CB-mKate2 plasmids. 48 hours post-electroporation, we added media supplemented with 20 μM A1155 to induce synchronous release of Shh and CD63. 90 minutes post release, we fixed cells as described above. We acquired Z-stacks through PLMCs using the super resolution (SR) mode on the LSM 980 with Airyscan2 (Zeiss) laser scanning confocal and produced maximum intensity projections using FIJI. We analyzed colocalization using the BIOP-JACoP plugin (https://github.com/BIOP/ijp-jacop-b), using the Otsu auto threshold to select Shh-CB-msfGFP and CD63-CB-mKate2 positive multivesicular bodies and Pearson’s Coefficients were calculated to quantify the level of overlap between the channels.

### Analysis of Shh Colocalization with Secreted Exosomes

To analyze what fraction of Shh colocalizes with secreted exosomes, we electroporated PLMCs with APX1-TetOn-3g, BITREX-CD63-pHluo-pm-mIRFP682, and APX2-Shh-FL-mScarlet2i plasmids, and then acquired data at 48 hours post electroporation.

To examine colocalization in the extracellular space, we acquired Z-stacks using the Marianas SoRa spinning disk confocal with a 100x objective with additional 2.8x magnification. We constructed max intensity Z-projections and constructed a cell body mask by thresholding the pm-IRFP682 membrane channel using FIJI’s Triangle threshold and Fill Holes function, and inverted this mask to create an ROI representing extracellular space. To detect and quantify colocalization between CD63-pHluo and Shh-FL-mScarlet2i puncta in the extracellular space ROI, we used the ComDet v.05.5 plugin for ImageJ (https://github.com/UU-cellbiology/ComDet) with the following parameters for both channels: “Include larger particles: true, Segment larger particles: true, Approximate particle size: 9, Intensity threshold (in SD): 40” and “Max distance between colocalized spots = 9”.

To examine colocalization along cytonemes, we acquired and projected Z-stacks as above, but without 2.8x additional magnification. We traced individual cytonemes visible within each projection manually to produce a composite ROI. We performed detection and co-localization of CD63-pHluo and Shh-FL-mScarlet2i puncta using the ComDet v.05.5 plugin the following parameters for both channels: “Include larger particles: true, Segment larger particles: true, Approximate particle size: 3, Intensity threshold (in SD): 40” and “Max distance between colocalized spots = 3”.

For both extracellular and cytoneme-associated puncta, we computed and report the fraction of exosome-bound Shh (CD63-pHluo+, Shh-FL-mScarlet2i+) divided by the total number of Shh-FL-mScarlet2i+ puncta.

### Extracellular (EC) Immunofluorescence staining

We generated PLMCs as described from the anterior half of limb buds (to ensure no endogenous Shh expression) and electroporated them with APX1-TetOn-3g and BITREX-Shh-FL-msfGFP-pm-mIRFP682 plasmids. 48 hours post electroporation, we incubated cells in a primary staining solution comprised of 5E1 anti-SHH primary antibody (1:50) in DPBS (Gibco) supplemented with 10% SuperBlock Blocking Buffer (Thermo Scientific) on ice for 25 minutes to prevent endocytosis. We rinsed cells 3x with ice cold DPBS and then incubated them with Fidelis Fix on ice for 15 minutes, followed by 3 washes with ice cold DPBS (5 mins each). We then incubated cells with FluoTag-X2 anti-Mouse IgG1 secondary antibody conjugated to Abberior STAR 580 dye (1:500, NanoTag Biotechnologies) in a 20% SuperBlock PBS solution for one hour on ice, followed by two 1-minute rinses and three 10-minute washes with ice cold PBS.

We acquired maximum intensity projected Z-stacks from EC immuno-stained cells using a 100x objective on the Marianas SoRa spinning disk confocal. To correct for non-specific background, we measured the average fluorescence intensity of control cells lacking primary antibody staining and subtracted this background from all images before analysis. We traced all individual cytonemes using the pm-mIRFP682 signal to produce a composite ROI, performed spot-based colocalization using the ComDet v.05.5 plugin with the same parameters listed above, and computed the fraction of extracellular Shh (Shh-FL-msfGFP+, Abberior STAR 580+) puncta divided by the total number of Shh-FL-msfGFP puncta for each cell.

### Quantifying MVB Fusion Events in PLMCs

To determine the effect of nexinhib20 incubation on MVB fusion/exosome secretion, we electroporated PLMCs with APX1-TetOn3g and APX2-CD63-pHluo plasmids, cultured them at 37°C for 24 hours, and then incubated the PLMCs with media containing both 1 μg/mL doxycycline and either 2 μM Nexinhib20 (MedChemExpress) or DMSO for another 24 hours. We imaged MVB fusion in PLMCs at room temperature (∼22°C) across both conditions using TIRF microscopy for 3 minutes at 300ms intervals with identical TIRF angle, laser power, and exposure. To correct for bleaching of the CD63-pHluo signal, we performed bleach Correction (Simple Ratio) using FIJI. We then generated cell body masks by creating an average intensity projection of the timelapse and thresholding the image using FIJI’s Triangle Auto Threshold.

To detect and quantify MVB fusion events in our cell mask ROIs, we used the ExoJ plugin in FIJI with the following parameters: Vesicle Detection include “Min Fusion event apparent size = 4 pixels (0.636 microns)”, “Max Fusion event apparent size = 10 pixels (1.589 microns), “Detection threshold = 18 sigma (wavelet scale)”, and the “Use wavelet filter” box checked. Parameters used for Spot Tracking (vesicle trajectory) include “Spatial searching range = 2 (0.318 microns)”, “Temporal searching depth (Gap closing) = 1 (frames)”, and “Minimal event size = 3 (frames).” Parameters used for Event Sorting and Analysis include “Min. points for fitting procedure = 5 (points)”, “Expanding frames (pre/post peak) = 4, 4 with Fixed position box checked”, “Detection threshold = 3 dF/σ (intensity)”, “Upper Decay limit = 20 frames (6s)”, “Lower Decay limit = 1 frame (0.3s)”, “Min. R^2^ (Decay fitting procedure) = 0.8”, “Min .R^2^ (Gaussian fitting procedure) = 0.85.” We calculated and reported the number of MVB fusion events per minute for each cell.

### Synthetic Shh Activity Gradient Formation and Analysis

To quantify how exosome and cytoneme perturbations affect Shh activity gradients in a tissue, we adapted a 2D synthetic reconstitution approach using a confluent field of Shh-Reporter “Receiver” cells and sparsely seeded Shh-expressing “Sender” cells.^15,48^ The “Receiver” cells were comprised of a stable NIH-3T3 cell line expressing a Shh activity Reporter (Gli Binding Site driving H2B-mCitrine) and were seeded on polymer coverslip bottom 24 well plate at a density of 4 x 10^5^ to act as a confluent seed layer to represent a mesenchymal tissue. The “Sender” cells were PLMCs collected exclusively from the anterior half of limb buds (free of endogenous Shh expression), electroporated with APX1-TetOn3g and BITREX-Shh-FL-msfGFP-pm-(mKate2 or mIRFP682), and sparsely seeded (500 cells per well) on top of the “Receiver” tissue. We then incubated each co-culture tissue with cell media supplemented with 2 μg/mL doxycycline for 72 hours followed by fixation as previously described.

To image the Shh signaling gradients in each well, we used the SlideBook acquisition software to image the entire mCitrine reporter gradient surrounding a membrane labeled (pm-mKate2 or pm-mIRFP682) Shh expressing PLMC using a two-color montage acquisition comprised of four separate 20 μm z-stacks (step size = 1.2 μm) centering around the Sender cell. We used identical laser power, exposure time, and gain in each image to ensure quantitative comparison For quantitative gradient analysis, we used FIJI software to open and create maximum intensity projections of all 4 z-stacks per gradient for both the Receiver cell (H2B-mCitrine) and the Sender cell membrane label (pm-mKate2 or pm-mIRFP682) channels. We stitched the 4 max projected images of each gradient, created a cell mask ROI around the membrane-labeled Sender cell, and measured the integrated density of the palmitoylated fluorescent protein membrane label (mKate2 or mIRFP682). As the palmitoylated fluorescent protein membrane label and Shh-FL-msfGFP are genetically coupled on the same plasmid, we used the integrated density of the membrane label acted as a proxy to normalize for each Sender cell’s level of Shh expression. Next, we used the StarDist 2D plugin in FIJI to segment and create nuclear ROIs in the H2B-mCitrine stitched image.^75^ Next, we calculated the center position of each gradient or “gradient centroid” positions by creating a rectangle ROI that exclusively highlighted the visible H2B-mCitrine gradient, measured the center of mass, bounding rectangle, and mean fluorescence of all nuclear ROIs within this gradient ROI, and then calculated the center of mass using nuclear position and mean values as weights. Using FIJI, we then created a 200 μm radius circle ROI centered around the gradient centroid coordinates and used the Radial Angle Profile plugin (http://questpharma.u-strasbg.fr/html/radial-profile-ext.html) to produce a profile plot of normalized integrated intensities around concentric circles as function of distance from the gradient centroid to the circle ROI boundary.

Considering each PLMC sender cell had variable amounts of Shh expression, we normalized the H2B-mCitrine profile plot values by dividing by the integrated intensity of the PLMC membrane fluorescent protein which again act as a proxy for Shh expression.

### Quantification and Statistical Analysis

We performed all statistical analysis using GraphPad Prism. Significance for comparisons between two experimental groups was determined with the Mann-Whitney U-test (2-tailed). Significance for comparison between more than two groups was determined by One-Way ANOVA with Tukey’s multiple comparison tests. ∗= p ≤ 0.05; ∗∗= p ≤0.01; ∗∗∗= p≤0.001, ∗∗∗∗=p<0.0001.

