## Supplemental Figures for "Exosome secretion is required for sonic hedgehog dispersal and signal gradient formation in the embryonic limb mesenchyme"

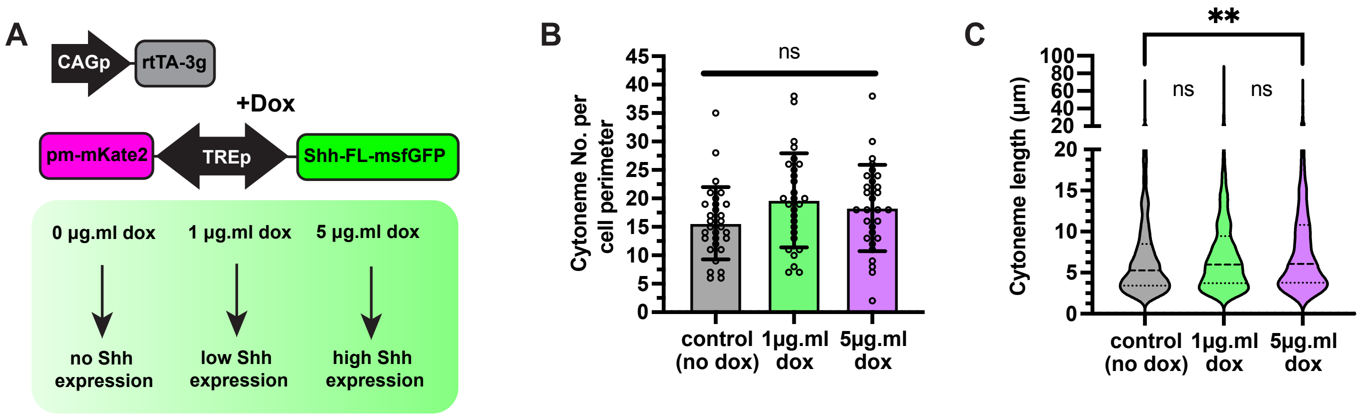

A
CAGp
rtTA-3g
+Dox
pm-mKate2
TREp
Shh-FL-msfGFP
0 µg.ml dox
1 µg.ml dox
5 µg.ml dox
no Shh expression
low Shh expression
high Shh expression
B
Cytoneme No. per cell perimeter
ns
control (no dox)
1µg.ml dox
5µg.ml dox
C
Cytoneme length (µm)
**
ns
ns
control (no dox)
1µg.ml dox
5µg.ml dox

**Supplemental Figure 1: Exogenous Shh expression has little to no effect on cytoneme production in PLMCs**

A) Diagram depicting experimental and genetic strategy to overexpress Shh in anterior limb bud derived PLMCs (no endogenous Shh expression). PLMCs were electroporated with a plasmid encoding TetOn-3g and a bidirectional Tet Response Element (TRE) promoter that drives expression of Shh-FL-msfGFP and pm-mKate2. To modulate expression of Shh, either 0, 1, or 5 mg.ml of Doxycycline was added to media to promote no, low, or high Shh overexpression, respectively. B) Quantification of average number of cytonemes per cell perimeter in all three overexpression conditions ($n = \sim 30$ cells per condition, +/- SD. One-way ANOVA performed ($p > 0.05$). C) Quantification of average cytoneme length per condition ($n > 490$ cytonemes per condition, +/-SD). One way ANOVA performed (** corresponds to $p = 0.099$).

**A**

| CB Isoform | Binding Affinity ($K_d$) | Release? |
|---|---|---|
| CB | 4 pM | NO |
| CB (E22A) | 270 pM | NO |
| CB (D138A) | 22 nM | YES |

(Shui et al., 2021)

**B**

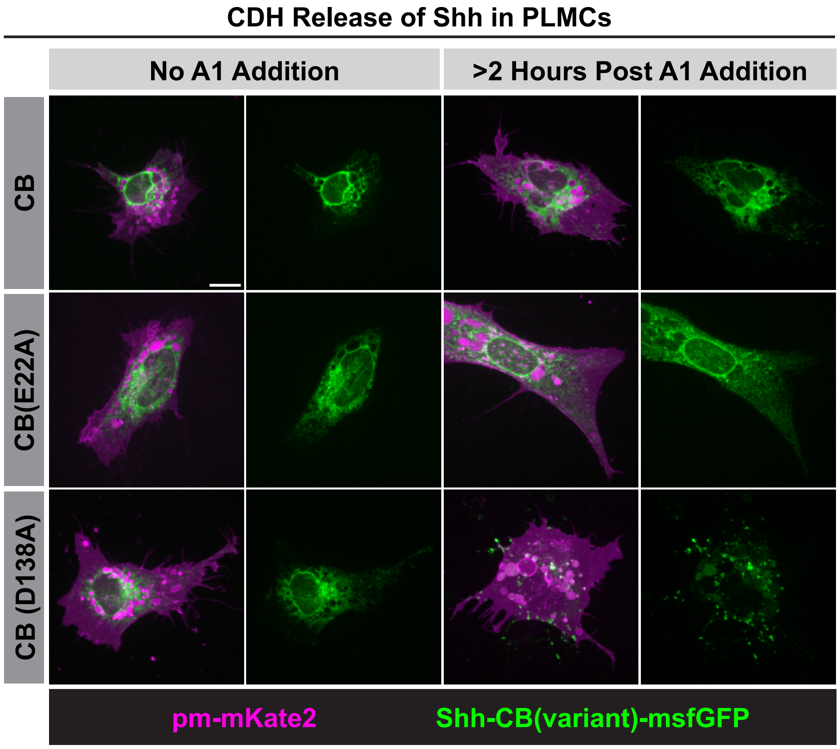


**C**

| | RUSH | CDH |
|---|---|---|
| Trap/Hook | Streptavidin | Bcl-xL |
| Binding Moiety | Streptavidin Binding Protein (SBP) | CDH Binding Moiety (CB) |
| Release Factor | Biotin | A-1155463 |
| Trap *In Vivo* | NO | YES |

**D**

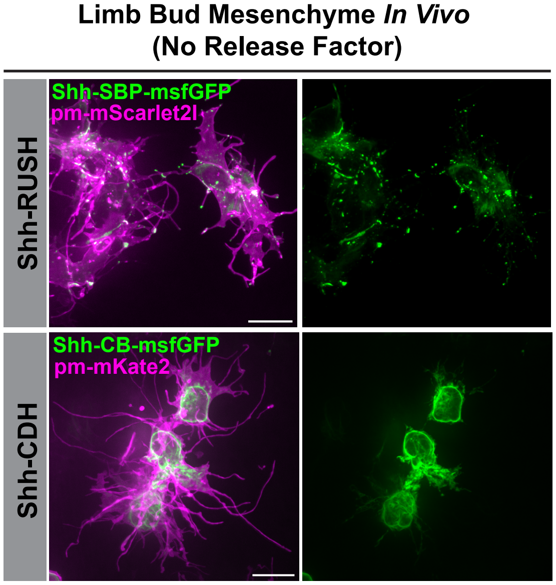

**Supplemental Figure 2: CDH release system provides tight ER trapping *in vitro* and *in vivo***
A) Table depicting previously published point mutated variants of the CB moiety and their binding affinity with Bcl-xL,[38] and if they are able to be rapidly released (under 2 hours) from an ER based trap in PLMCs. B) Micrographs showing PLMCs that were electroporated with the ER-localized Bcl-xL trap and a Shh-CB(variant)-msfGFP with and without ~2 hours of A1 incubation. Note that the CB and CB(E22A) variants, that have higher affinity with Bcl-xL, do not release from the ER despite 2 hours of incubation with the A1 release factor. However, the CB(D138A) variant can both trap in the ER and release upon A1 incubation. C) Table depicting a comparison between RUSH and CDH synchronous release systems. D) Micrographs depicting limb bud mesenchymal cells, *in vivo,* either electroporated with RUSH (ER-localized streptavidin and Shh-SBP-msfGFP) or CDH (ER-localized Bcl-xL and Shh-CB-msfGFP) release systems without any addition of release factors (Biotin or A1, respectively). Note that Shh-SBP-msfGFP (RUSH) is not trapped in the ER and exists as punctate structures throughout the cell body and cytonemes. Whereas Shh-CB-msfGFP (CDH) remains tightly trapped in the ER.

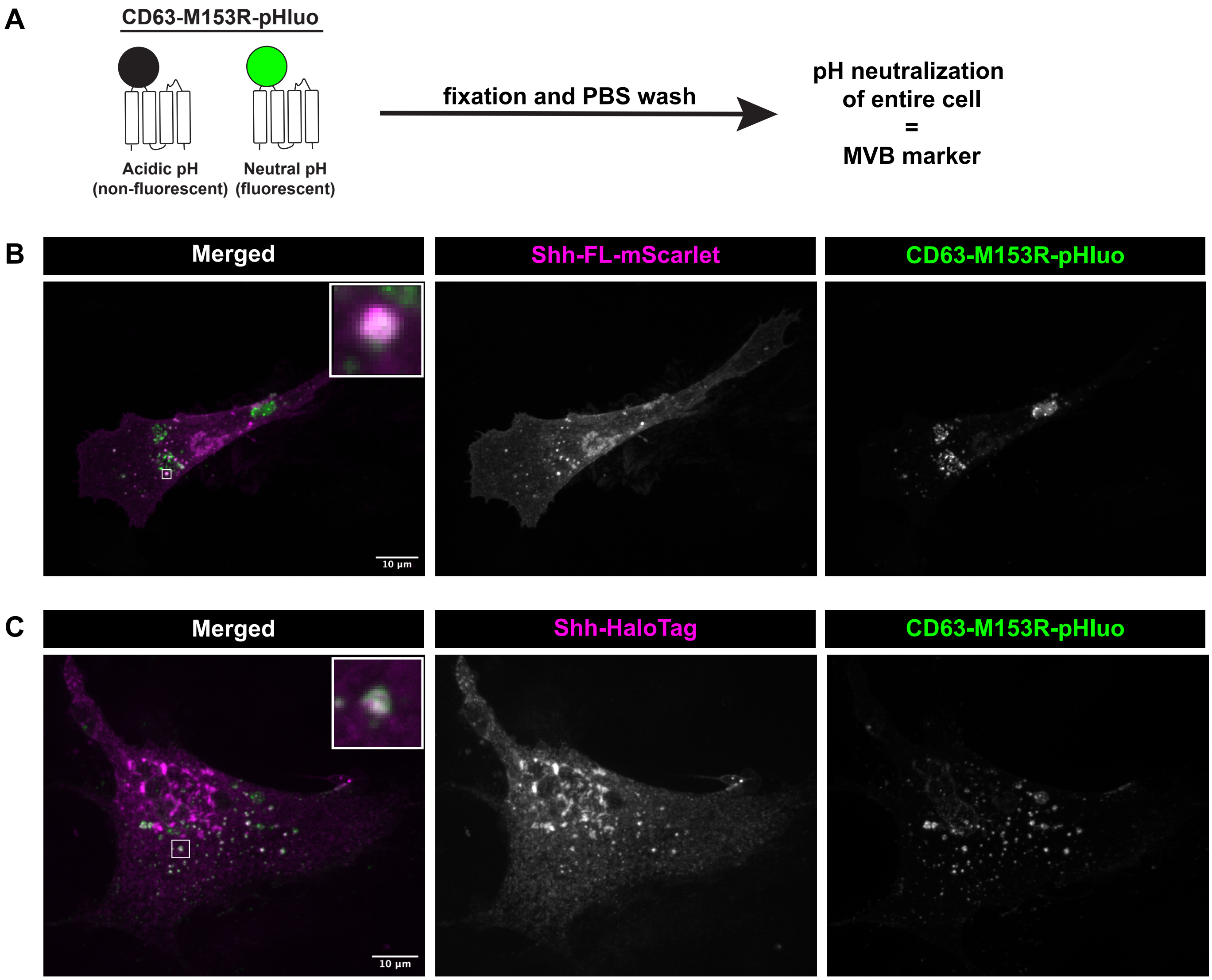
A
CD63-M153R-pHluo
Acidic pH
(non-fluorescent)
Neutral pH
(fluorescent)
fixation and PBS wash
pH neutralization
of entire cell
=
MVB marker
B
Merged
Shh-FL-mScarlet
CD63-M153R-pHluo
10 µm
C
Merged
Shh-HaloTag
CD63-M153R-pHluo
10 µm

**Supplemental Figure 3: Multiple Shh fusion protein variants accumulate in MVBs in PLMCs**
A) Schematic showing that CD63-pHluo in fixed cells acts as an MVB/exosome marker. B) Micrographs of PLMCs co-expressing Shh-FL-mScarlet and CD63-pHluo. Note high degree of colocalization in endosome-like structures (inset). C) Micrographs of PLMCs co-expressing Shh-HaloTag (+ Janelia Fluor JFX650) and CD63-pHluo. Again, note high degree of colocalization in endosome-like structures (inset).

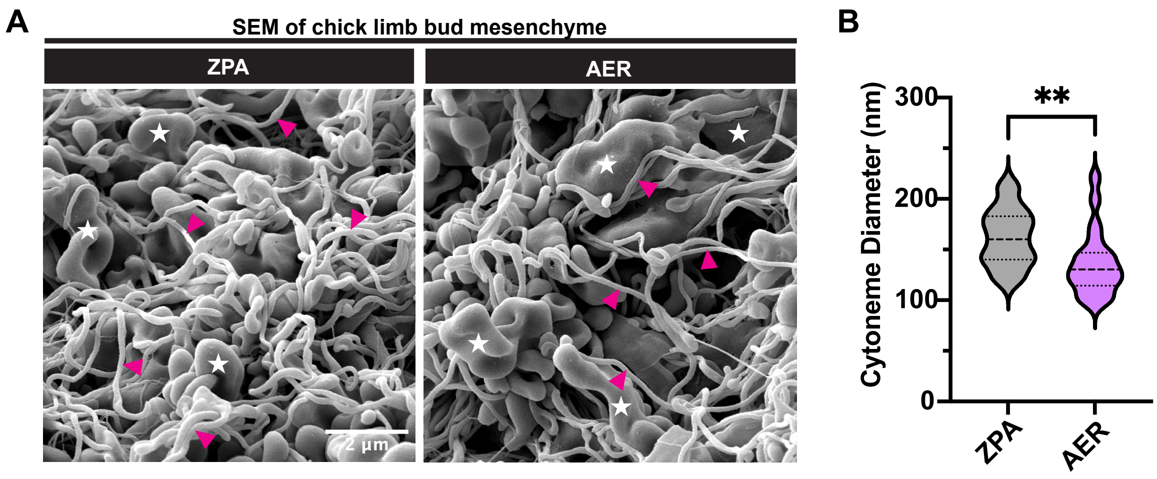
A
SEM of chick limb bud mesenchyme
ZPA
AER
2 µm
B
**
Cytoneme Diameter (nm)
300
200
100
0
ZPA
AER

**Supplemental Figure 4: Ultrastructural views of limb bud mesenchyme in vivo show vast size discrepancies between cytoneme and MVB diameters**

A) Scanning electron micrographs of surface limb bud mesenchyme (ectoderm removed) from HH 22-24 chick embryos. Left panel is an example micrograph of the ZPA, while the right is an example micrograph of the mesenchyme underlying the AER region at the distal tip of the limb bud. Note globular cell bodies (white stars) and complex network of cytonemes (magenta arrowheads). B) Graph represents the composite (average) diameter of cytonemes in the ZPA and AER. Although the cytonemes are slightly wider in the ZPA, they still are under 200 nm in diameter.

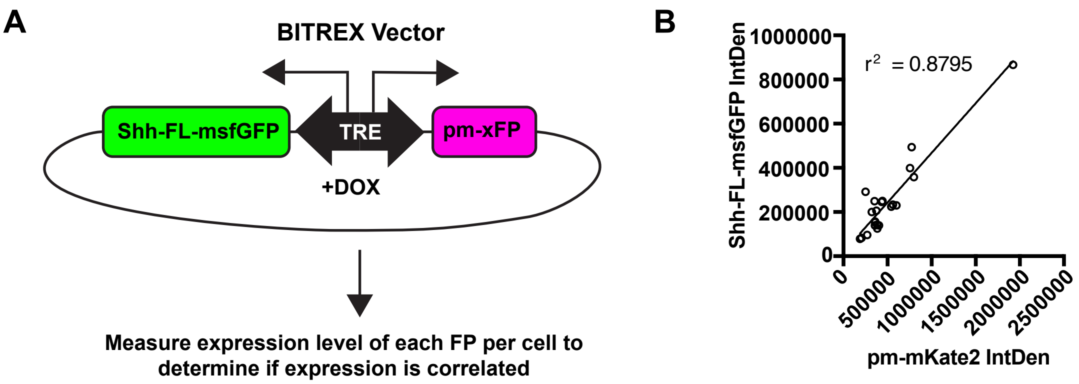
A
BITREX Vector
Shh-FL-msfGFP
TRE
pm-xFP
+DOX
Measure expression level of each FP per cell to determine if expression is correlated
B
$r^2$ = 0.8795
Shh-FL-msfGFP IntDen
1000000
800000
600000
400000
200000
0
0
500000
1000000
1500000
2000000
2500000
pm-mKate2 IntDen

**Supplemental Figure 5: BITREX vectors provide genetically coupled expression of downstream constructs**

A) Schematic depicting map of BITREX vector. Bidirectional Tet Response Element (TRE) promoters drive expression of both Shh-FL-msfGFP and pm-mKate2 or pm-mIRFP682. With both TetON3g expressed and doxycycline added, both constructs are expressed and their relative expressions can be measured in PLMCs via Fluorescence Intensity. B) Graph depicting the correlation of integrated density of Shh-FL-msfGFP and pm-mKate2 per cell (n = 22 cells). A linear regression was performed with $r^2 = 0.8795$, suggesting tight correlation.

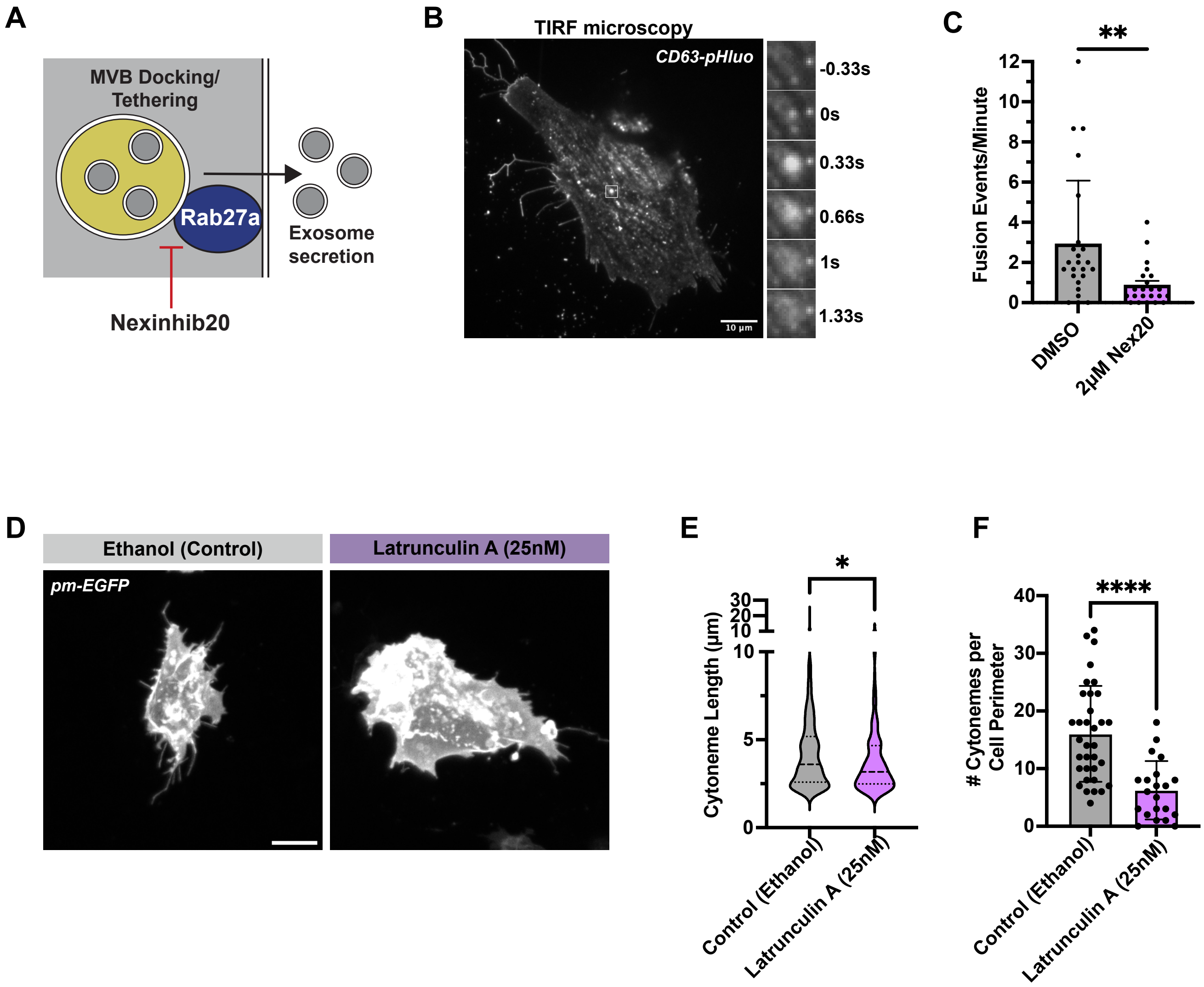
A
MVB Docking/
Tethering
Rab27a
Exosome
secretion
Nexinhib20
B
TIRF microscopy
CD63-pHluo
10 µm
-0.33s
0s
0.33s
0.66s
1s
1.33s
C
**
Fusion Events/Minute
DMSO
2µM Nex20
D
Ethanol (Control)
Latrunculin A (25nM)
pm-EGFP
E
*
Cytoneme Length (µm)
Control (Ethanol)
Latrunculin A (25nM)
F
****
### Cytonemes per
Cell Perimeter
Control (Ethanol)
Latrunculin A (25nM)

**Supplemental Figure 6: Small molecule inhibitors of Rab27a-JFC1 interaction and of actin polymerization can perturb MVB fusion and cytoneme production in PLMCs**

A) Diagram depicting Rab27a inhibition via the small molecule inhibitor Nexinhib20 (Nex20) and its effect on MVB docking/fusion. B) Micrograph of PLMC expressing CD63-pHluo. We used total internal reflection fluorescence (TIRF) microscopy to image MVB fusion events characterized by large flashes of CD63-pHluo expression at the PM followed by exponential decay of signal. One MVB fusion event (magnified white box) is displayed on right of image. C) Quantification of MVB fusion events in PLMCs incubated with either DMSO or Nex20 (2 μM) for 24 hours. Quantification was performed using ExoJ plugin in FIJI (n >22 cells, Mann-Whitney U test $p = 0.0019$). D) Example micrographs of PLMCs expressing pm-EGFP to visualize cytonemes after Ethanol or Latrunculin A (LatA) (25nM) treatment (24hrs) – Scale Bar = 10μm. E) Quantification of cytoneme lengths of in Ethanol and LatA treated PLMCs (n = 539 cytonemes in Ethanol treatment, n = 131 cytonemes in LatA treatment, Mann-Whitney U test $p = 0.0185$). F) Quantification of average cytoneme number per cell perimeter in ethanol and LatA treated PLMCs (n = 34 cells in Ethanol treatment, n = 21 cells in LatA treatment, Mann-Whitney U test $p < 0.0001$).
